# FAMetA: a mass isotopologue-based tool for the comprehensive analysis of fatty acid metabolism

**DOI:** 10.1101/2022.05.11.491462

**Authors:** María Isabel Alcoriza-Balaguer, Juan Carlos García-Cañaveras, Marta Benet, Oscar Juan Vidal, Agustín Lahoz

**Author notes:** These authors contributed equally (María Isabel Alcoriza-Balaguer, Juan Carlos García-Cañaveras).

## Abstract

The use of stable isotope tracers and mass spectrometry (MS) is the gold standard method for the analysis of fatty acids (FAs) metabolism. Yet current state-of-the-art tools provide limited and difficult to interpret information about FA biosynthetic routes. Here we present FAMetA, an R-package and a web-based application (www.fameta.es) that use ^13^C mass-isotopologue profiles to estimate FA import, de novo lipogenesis, elongation, and desaturation in a user-friendly platform. The FAMetA workflow covers all the functionalities needed for MS data analyses. To illustrate its utility, different *in vitro* and *in vivo* experimental settings are used in which FA metabolism is modified. Thanks to the comprehensive characterisation of FA biosynthesis and the easy-to-interpret graphical representations compared to previous tools, FAMetA discloses unnoticed insights into how cells reprogramme their FA metabolism and, when combined with FASN, SCD1 and FADS2 inhibitors, it enables the straightforward identification of new FAs by the metabolic reconstruction of their synthesis route.

## Introduction

Fatty acids (FAs) are key metabolites that play a central role in cellular biology. FAs act as building blocks for the synthesis of complex lipids or as a source of energy, but also as signalling molecules ^1^. Dysregulated FA metabolism has been associated with many of the most prevalent diseases, including obesity ^2^, type 2 diabetes ^3^, non-alcoholic fatty liver disease ^4^ or cancer ^5^. FAs can be either synthesised *de novo* inside cells or imported from external sources. The main product of *de novo* lipogenesis (DNL) is palmitic acid [FA(16:0)], which results from the condensation of acetyl-CoA molecules through the enzymatic action of acetyl-CoA carboxylase (ACACA/B) and FA synthase (FASN). The acetyl-CoA pool is generated via ATP citrate lyase (ACLY) from citrate which can, in turn, be produced from several carbon sources (i.e., glucose, glutamine, amino acids, FAs), or from acetate *via* acetyl-CoA synthetases (ACSS1/2) ^6^. Linoleic [FA(18:2n6)] and α-linolenic acid [FA(18:3n3)] are essential FAs that must be exogenously acquired. Free FA import occurs by either passive diffusion or the action of translocases like CD36 and FA transport proteins (FATPs). FAs can be elongated via the elongation of very long-chain FAs proteins (ELOVL1-7). They can also be desaturated via the action of stearoyl-CoA desaturases 1/5 (SCD1/5) and FA desaturases 1/2 (FADS1/2) enzymes ^1,5^. The wide variety of FAs required for the cellular functioning results from these transformations.

Stable-isotope tracing combined with mass spectrometry (MS)-based detection is a widespread method for interrogating FA metabolism. The total FA synthesis rate can be estimated by using D_2_O, which labels FAs through direct solvent incorporation and NADPH-mediated hydrogen transfer ^7,8^. Additionally, employing ^13^C-labelled tracer nutrients (e.g. U-^13^C-glucose, U-^13^C-glutamine, U-^13^C-acetate, etc.) allows the total FA synthesis rate and the relative contribution of a given nutrient to be estimated ^9^. The framework for FA synthesis data analysis using ^13^C-labelled tracers and MS was initially set up by Isotopomer Spectral Analysis (ISA) ^10^ and Mass Isotopomer Distribution Analysis (MIDA) ^11^, which model FAs synthesis following the incorporation of n 2-carbon units using multinomial distribution fitting. Unfortunately, these mass isotopologue modelling methods only provide information about the DNL of FAs for which the contribution of elongation is minimal (i.e., FAs of 14 or 16 carbons) ^10–12^. ConvISA incorporated one elongation step to, thus, extending the analysis to 18-carbon FAs ^13^. Recently, Fatty Acid Source Analysis (FASA) included many elongation steps, which extend the FAs species that can be properly modelled to 26 carbons ^14^. FASA has some limitations because it assumes *de novo* synthesis up to 26- carbon FAs and it calculates multiple import-elongation terms (i.e. *IE*_*n*_, which refers to imported and elongated *n* times), which does not accurately represent the actual biological process. A simple strategy for estimating the desaturation of FA(18:0) to FA(18:1n9) has been described ^15^. However, this approach is based on the total labelling of precursor and product FAs ^15^, and its application to the complete array of desaturations has not yet been explored. Despite these valuable advances, reliable FA elongation calculations are still to be fully addressed, whereas systematic desaturation estimations remain unresolved. Additionally, the above-mentioned algorithms are developed for platforms that require computational skills and commercial software, thus, they are not readily accessible to the broad metabolism community. To bridge this gap, we developed FAMetA (Fatty Acid Metabolism Analysis), a mass isotopologue-based tool implemented as R-package and web-based application that aims to analyse all the biosynthetic reactions within the FA metabolic network. FAMetA provides all the functionalities needed to analyse MS data and returns easy-to-interpret results that facilitate straightforward FA metabolism analyses and the identification of unknown FAs.

## Results

### FAMetA overview

FAMetA is an R package (https://CRAN.R-project.org/package=FAMetA) and a web-based platform (https://www.fameta.es) that rely on mass isotopologue distributions from GC-MS or LC-MS to estimate the import (*I*), the synthesis of FA(14:0)/FA(16:0) (*S*), the fractional contribution of the ^13^C- tracer (*D*_*0*_, *D*_*1*_, *D*_*2*_, which represent the acetyl-CoA fraction with 0, 1 or 2 atoms of ^13^C, respectively), the elongation (*E*) and the desaturation (*Δ*) parameters for the expected network of FA synthesis reactions up to 26-carbons ^16^ (**Figure 1, Extended Data Figure 1**). To accurately determine these parameters, the analytical method of choice must allow the detection of both the multiple isotopologues associated with a given FA and the chromatographic resolution of isomers (e.g. 18:1n7, 18:1n9, 18:1n10, 18:1n7trans). To this end, we developed a derivatisation-free C18-UPLC method that provides the required separation of key biologically relevant isomers (e.g. 16:1n5/7, 16:1n9, and 16:1n10 or 18:1n7, 18:1n9, 18:1n10 and 18:1n7trans), and enables their sensitive detection by high-resolution MS (HRMS) (**Supplementary Figure 1**). The FAMetA workflow comprises all the functionalities needed, from data preprocessing to group-based comparisons and graphical output (**Figure 1, Supplementary Figures 2-3**). Mass isotopologue distributions usually show overdispersion, which can be attributed to the cellular heterogeneity, time-dependent variations that result from changes in nutrient availability, differences between the various intracellular FA pools (e.g. differences between lipid classes or between FA/lipids located in different organelles), among others. FAMetA implements quasi-multinomial modelling that improves the fitting of mass isotopologue distributions compared to formerly used multinomial modelling^10–14,17,18^. Furthermore this fitting provides the parameter *Φ* that accounts for data overdispersion (**Extended Data Figure 2**). For FAs up to 16 carbons, the DNL parameters (*I, S, Φ* and D_0_, D_1_, D_2_) are estimated. The equations employed to fit the experimental isotopologue distribution are equivalent to those employed by the ISA algorithm^11,12^ if the parameter *Φ* = 0 and if the ISA equations are modified to take into account that the data has been corrected for the natural abundance of ^13^C. For the FAs of 18 to 26 carbons, apart from the parameters *S* and *I*, up to five elongation terms (*E*_*n*_, n=1 for 18-carbon to n=5 for 26-carbon FAs) are estimated. Each elongation term represents the direct estimation of the fraction that comes from the elongation of the total pool of the precursor FA (**Extended Data Figure 3**). Compared to previous tools (i.e. FASA, where the synthesis of a FA longer than 16C is described as DNL up to the total length and multiple import-elongation terms are implemented^14^), the way in which elongations are calculated by FAMetA leads to more comprehensible elongation parameters and to a more intuitive and straightforward biological interpretation of the results. For the FAs that result from the direct desaturation of one precursor FA, *Δ* is indirectly estimated based on the calculated synthesis parameters of the precursor (*S* or *E*) and the FA of interest (*S’* or *E’*) (i.e. *Δ*= *S’/S or Δ*= *E’/E*) (**Extended Data Figure 3**; see **Online Methods** for further details). The strategy proposed here is inspired by the simple approach described by the previous work developed by Kamphorst et *al* ^15,19^. The authors calculate desaturation for FA(18:1n9) based on the total labelling in FA(18:0) and FA(18:1n9). We extend the strategy to the complete set of desaturations within the FA metabolic network and refine the calculation by using an approach that uses the estimated synthesis parameter of interest instead of total labelling. The use of the complete isotopologue distribution to estimate the substrate and product FA synthesis parameters of interest instead of a single summed value may lead to a more robust and accurate estimation of desaturation. However, in our opinion the key advance of FAMetA is about the possibility of estimating alterations in concrete desaturation steps from the complete set of parameters calculated for a given FA (e.g. identify alterations in SCD activity between two conditions based on the information obtained for FA(18:1n7), where the double bond is introduced at the 16-carbon level). Finally, the complete metabolic network of FA synthesis is summarised for each sample and group, and comparisons between groups are made and graphically represented (**Figure 1**). As in previous tools (i.e. ISA, ConvISA and FASA^11–14^) the de novo synthesis parameters (*S, E, Δ*) are time-dependent. Therefore, at any given time, such parameters correspond to the fraction of a particular FA that has been de novo synthesised up-to-the moment of the sampling, and it corresponds to the actual portion of FA that comes from DNS only if the steady-state has been achieved. Accordingly, the import term (*I*=1-*S* or *I*=1-*E*_*n*_) accounts for both import and pre-existing FAs at any given time and to the actual fraction that is acquired from the exogenous pool when the steady state has been reached. The conditions of metabolic and isotopic steady states are only achieved, or can be closely approximated, if the cells are cultured during a long-enough time to ensure that the pre-existing FA pools can be diluted out while ensuring a nutrient supply that maintains relatively stable concentrations^14,20^ A comparison of the functions implemented by FAMETA and other available tools is summarized in **Supplementary Table 1**.

**Figure 1.**
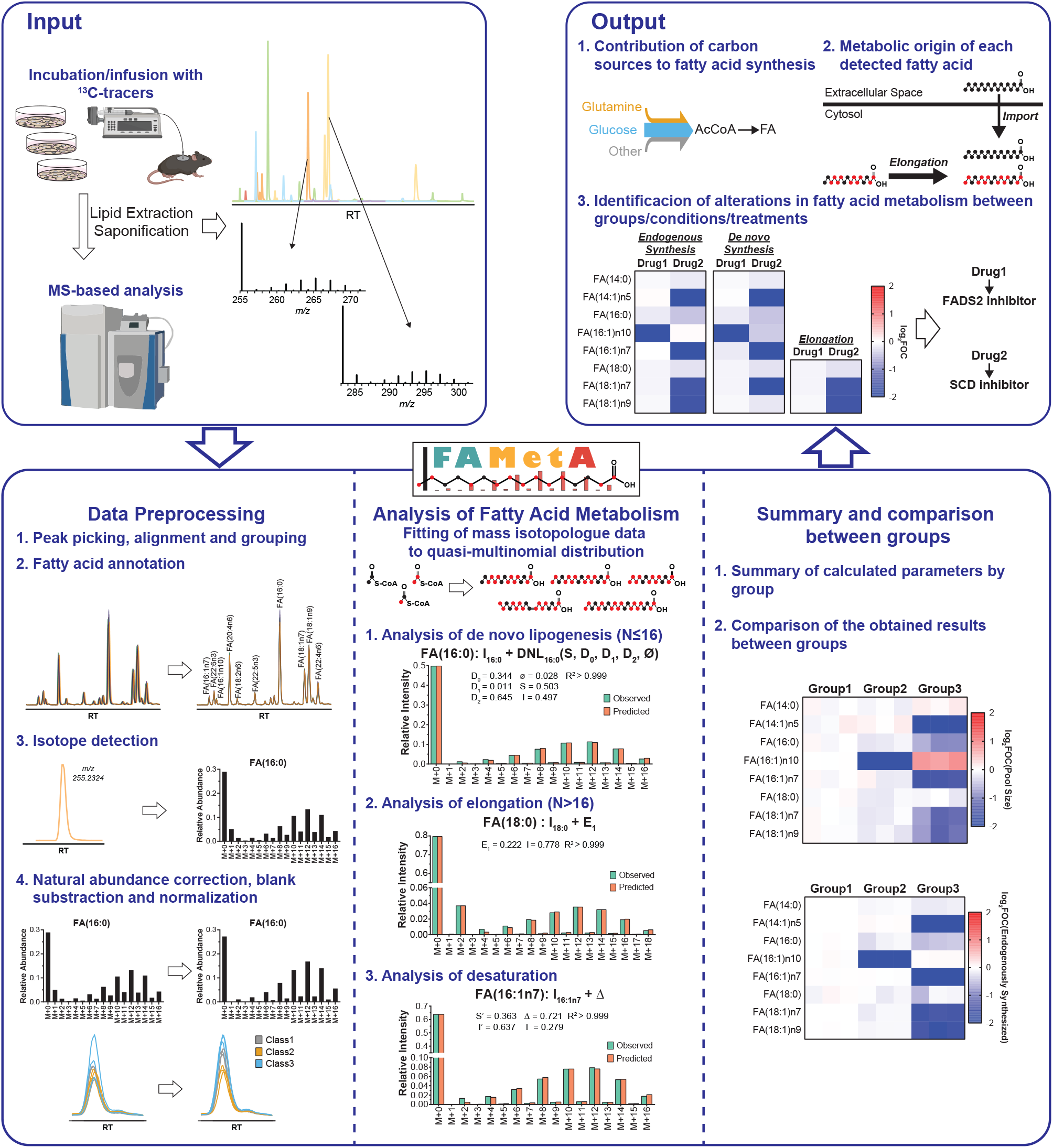
FAMetA workflow. FAMetA is an open-access platform-independent software for estimating FA metabolism based on mass isotopologue data that can be executed locally in an R environment (https://CRAN.R-project.org/package=FAMetA) or online (www.fameta.es). FAMetA uses as input the data generated after incubation/infusion with suitable ^13^C-tracers, based on the LC-MS or GC-MS analysis of FA extracts. Briefly, the FAMetA workflow comprises the following steps: 1) data preprocessing; 2) analysis of the FA metabolism for each sample and detected FA; 3) the combination of these individual results to provide an overview of the FA metabolism network for each condition of interest and to compare them. The analysis of the FA metabolism parameters is based on fitting the experimental mass isotopologue distribution to a quasi-multinomial distribution. For FAs of 14C and 16C, the imported fraction (*I*) and the DNL parameters (i.e. synthesis (*S*), fractional contribution of the tracer (*D*_*0*_, *D*_*1*_ and *D*_*2*_) and overdispersion (*Φ*) are calculated. For FAs of more than 16C, the *D*_*0*_, *D*_*1*_, *D*_*2*,_ and *Φ* parameters are imported and the sources are described as the import (*I*) plus the elongation (*E*_*n*_) of the total pool of the precursor FA [e.g. for FA(18:0), sources are described as *I*_*18:0*_ + *E*_*1*_= *I*_*18:0*_ + *E*_*1*_ (*I*_*16:0*_ + *S*)]. For the FAs that are the result of desaturation, sources are described as the import (*I*), plus desaturation (*Δ*), where *Δ* is indirectly estimated according to the synthesis parameters of the precursor (*S* or *E*) and product FAs (*S’* or *E’*), where *Δ* = *S’*/*S* or *E’*/*E*. Depending on the experimental design, the most relevant biological outputs to be obtained include the fractional contribution of each tested carbon source, the detailed description of the metabolic origin of each detected FA, and the elucidation of an alteration in the FA metabolism between conditions of interest.

### FAMetA validation

*In silico* mass isotopologue distributions are generated to validate the FAMetA algorithm. To simulate experimental distributions, multiple values covering the expected range for each parameter are used. For each theoretical isotopologue distribution, 10 realisations of Gaussian noise are simulated at four noise levels [0%, 2%, 5%, or 10% relative standard deviation (RSD)]. The generated data are used to calculate the RSD and relative error of each modelled synthesis parameter for the following FAs, which comprise an example of all the reactions included in FAMetA: FA(16:0) (**Supplementary Figure 4**), FA (18:0) (**Supplementary Figure 5**), FA(20:0) (**Supplementary Figure 6**), FA(22:0) (**Supplementary Figure 7**), FA(24:0) (**Supplementary Figure 8**), FA(16:1n7) (**Supplementary Figure 9**), and FA(18:1n9) (**Supplementary Figure 9**). FAMetA accurately determines the complete set of FA synthesis parameters (relative error < 15%, RSD < 15%) whenever the fractional contribution of the tracer (*D*_*2*_) and the parameters to be calculated for a given FA (i.e. *S, E*_*1*_, *E*_*2*_, *E*_*3*_, and *E*_*4*_) fall within the 0.05 - 0.9 range. This ensures its applicability in an actual biological scenario.

### FAMetA enables straightforward FA metabolism analyses

To evaluate FAMetA performance, a variety of *in vitro* and *in vivo* experimental settings are used. Firstly, mouse CD8^+^ T-cells are incubated for 72 h with different uniformly ^13^C labelled tracers (U-^13^C-glucose, U-^13^C-glutamine, U-^13^C-lactate or U-^13^C- acetate) in the presence or absence of well-known inhibitors of FA metabolism enzymes [i.e. FASN (GSK2194069, FASNi) ^21^, SCD1 (A93572, SCD1i) ^22,23^ and FADS2 (SC26196, FADS2i) ^24^]. Total lipids are extracted from cell pellets and saponified to release FAs, which are subsequently analysed by our in-house LC-HRMS method. The FAMetA mass isotopologue data analysis satisfactorily estimates the FA metabolism parameters for the 27 known detected FAs (**Extended Data Figure 4, Supplementary Results**). Treatment with FASNi and SCD1i slightly decreases cell proliferation, but FADS2i does not (**Figure 2a**). Changes in the relative pool size of the detected FAs appear (**Figure 2b**); e.g. SCD1i lowers the intracellular levels of the n5, n7, and n9 series FAs, and increases the relative abundance of FADS2 products [e.g. sapienic acid, FA(16:1n10)], while FADS2i considerably diminishes sapienic acid abundance, which is consistent with previous reports on the complementary and compensatory roles of SCD1 and FADS2 ^25^ (**Figure 2b**). When analysing endogenous synthesis, the changes reveal which enzymes are involved in the synthesis of each identified FA. FASNi decreases the endogenous synthesis of all the FAs that come from FA(16:0), and SCD1i and FADS2i decreases the endogenous synthesis of all the FAs that these enzymes are involved in (e.g. n9 series FAs for SCD1i, n10 series FAs for FADS2i) (**Figure 2c**). When focusing on each calculated synthesis parameter, identifying the step in which each enzyme acts and mapping synthesis routes are straightforward. For example, for FA(18:1n7) and FA(18:1n9), SCDi differentially affects synthesis parameters. In FA(18:1n9), where SCD acts at the 18-carbon level, the most prominent decrease is in calculated *E*_*1*_ (i.e. *E*_*1*_’= *E*_*1*_\**Δ*), in FA(18:1n7), where SCD acts at the 16-carbon level, both calculated *S* (i.e. S’= *S*Δ*), and *E*_*1*_ decreases upon treatment with SCDi (**Figure 2d-e**). The SCDi inhibition pattern observed in FA(18:1n9) is mirrored in FA(20:1n9) and FA(20:3n9) (**Figure 2f-g**). In addition, FADS2i decreases the calculated *E*_*2*_ (i.e. *E*_*2*_’= *E*_*2*_\**Δ*) for FA(20:3n9), which is indicative of FADS2 introducing a double bond at the 20-carbon level (**Figure 2g**). Thus FAMetA allows the identification of both changes in general patterns and particular synthesis parameters induced by FA metabolism inhibitors.

**Figure 2.**
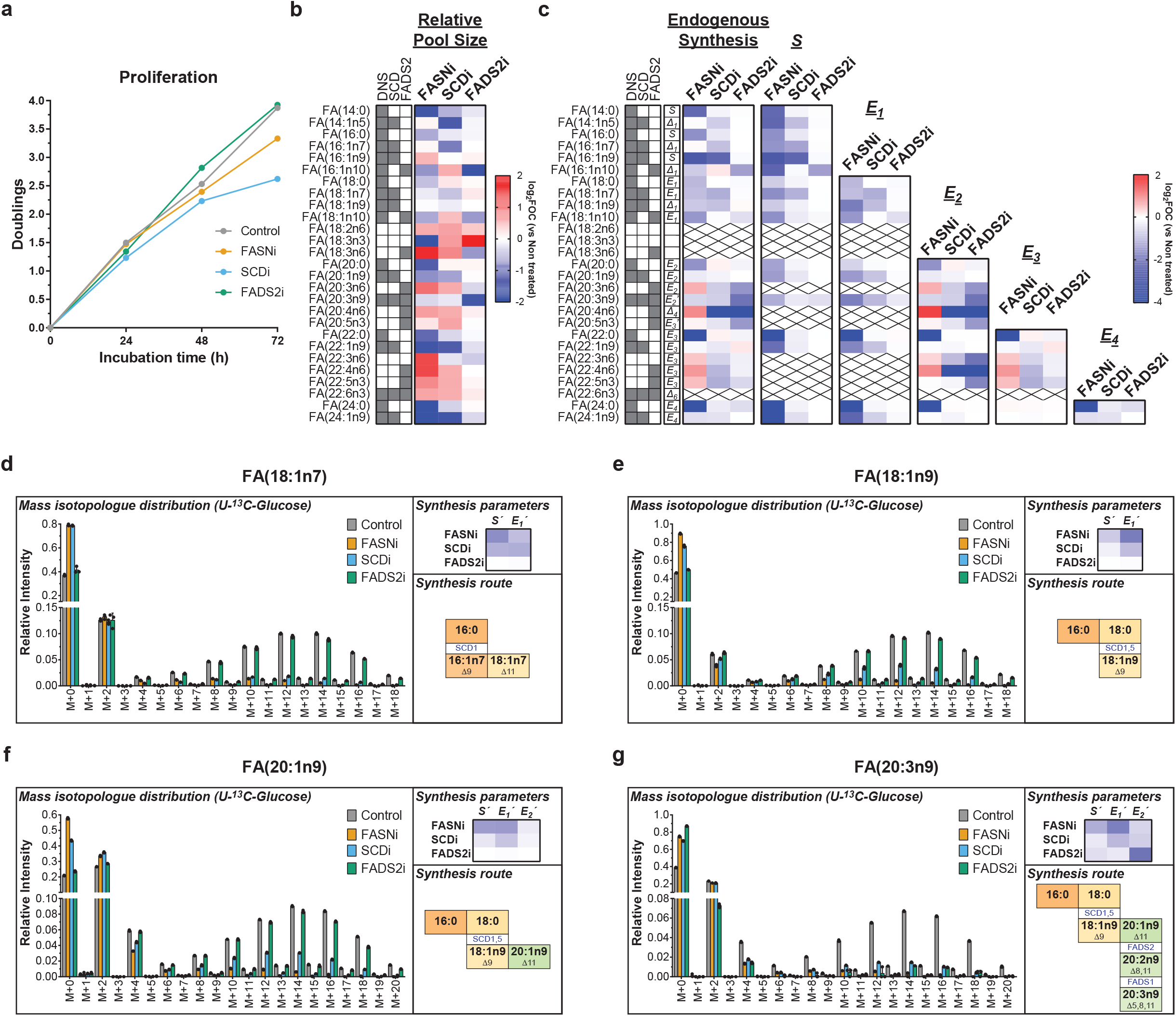
FAMetA enables the straightforward detection of alterations in the FA metabolic network. Analysis of alterations in FA biosynthesis in the active mouse CD8^+^ T-cells incubated for 72 h with ^13^C- glucose induced by FASN inhibitor GSK2194069, SCD inhibitor A93572 and FADS2 inhibitor SC26196. **a**, Mean proliferation of the active mouse CD8^+^ T cells during the 72-hour incubation period. **b**, Heatmap showing for each identified FA the mean value of the log_2_ fold-of-change (vs. untreated) in the relative pool size. **c**, Heatmap showing the mean value of the log_2_ fold-of-change (vs. untreated) for each identified FA in the following parameters: endogenously synthesised fraction, calculated *S, E*_*1*_, *E*_*2*_, *E*_*3*_ and *E*_*4*_. For each FA, the parameter reported for the endogenous synthesis is indicated. **d-g**, Mass isotopologue distribution, the mean value of the log_2_ fold-of-change (vs. untreated) in the synthesis parameters and synthesis route for FA(18:1n7) (**d**), FA(18:1n9) (**e**), FA(20:1n9) (**f**) and FA(20:3n9) (**g**). In all cases n=3. Individual points are shown for the mass isotopologue distributions, and the mean values are reported elsewhere. The shadowed cells in **b** and **c** indicate the activities (DNS, SCD or FADS2) involved in the synthesis of a particular FA. On the heatmaps, crosses indicate missing or NA values. In **d-g, the** horizontal transitions in the synthesis route description denote elongations (enzymes not indicated), and vertical transitions denote desaturations (enzymes indicated). See **Extended Data Figure 3** as a guide as to how calculations are done and which parameters are reported as the endogenous synthesis of each identified FA.

Then we move on to analyse previously published data on the H1299 cells incubated with U-^13^C-glucose and U-^13^C-glutamine, where the down-regulation of SREBP cleavage activating protein (SCAP), a key protein in the regulation of FA metabolism, is induced ^14^. SCAP down-regulation decreases the synthesis of monounsaturated n7 [i.e. FA(16:1n7) and (FA(18:1n7)] and n9 [i.e. FA(18:1n9), FA(20:1n9), and FA(22:1n9)] FAs (**Extended Data Figure 5**). When focusing on particular synthesis parameters, the calculated *S* (i.e. *S’*= *S*Δ*) is the most altered parameter for the n7 series, which is consistent with SCD1 introducing the double bond at the 16-carbon level. The calculated *E*_*1*_ (i.e. *E*_*1*_’= *E*_*1*_\**Δ*) is the most altered parameter for the n9 series, and is consistent with SCD1 introducing the double bond at the 18-carbon level (**Extended Data Figure 5b**). While the authors of the study conclude that SCAP down-regulation causes a decrease in both FASN activity and elongation ^14^, our refined analysis, which includes the calculation of desaturation and easy-to-interpret direct estimations of each elongation step, identifies that the main decrease occurs in the endogenous synthesis of the SCD1-derived n7 and n9 series of FAs. This indicates diminished SCD1 activity as the main metabolic change induced after SCAP silencing (see **Supplementary Information Results** for a detailed comparison of the performance of FAMetA and FASA).

Finally, we analyse previously published data on the incorporation of U-^13^C-fructose into saponified circulating FAs in wild-type and intestine-specific ketohexokinase (KHK-C) knockout mice after drinking normal water for 8 weeks, or 5% or 10% sucrose water ^26^. FAMetA properly fits the *in vivo* generated data, characterised by low synthesis, slight contribution and a high proportion of odd-labelled isotopologues, but within the high-confidence ranges established using the *in silico* validation datasets (**Extended Data Figure 6a-d**). The observed general trend suggests increased DNL, elongation and desaturation upon sucrose treatment, with a more pronounced effect on the KHK-C knockout mice (**Extended Data Figure 6e**). Neither sucrose nor KHK-C ablation influences the fractional contribution of fructose to DNL (**Extended Data Figure 6f-g**), but exposure to drinking fructose significantly alters DNL (S), elongation (E_1_ and E_2_) and desaturation (**Extended Data Figure 6h-k**). The *post hoc* comparisons reveal significantly heightened FA(16:0) synthesis when drinking more fructose, but only in the KHK-C knockout group (**Extended Data Figure 6h**), as well as augmented desaturation when drinking more fructose in both the wild-type and KHK-C knockout mice (**Extended Data Figure 6k**). These results agree with and extend those reported by the authors of the study by showing increased total ^13^C-labelled carbons in saponified circulating palmitate, which accounts for the cumulative effect of DNL, the contribution of fructose to DNL and the palmitate concentration^26^.

### FaMetA enables the identification of unknown FAs in biological samples

The analysis of total FAs in non-small cell lung cancer (NSCLC) cell line A549 reveals high FAs diversity (62 species), including several Fas (33) that do not match available standards (**Extended Data Figure7**). We hypothesise that the information provided by the retention time of each FA combined with the FAMetA analysis of the MS-data generated using U-^13^C-glucose and well-characterised inhibitors (i.e. FASNi, SCDi, and FADS2i) would provide a valuable strategy to identify unknown and unexpected FAs by the reconstruction of their metabolic synthesis route. All the detected unknown FAs incorporate ^13^C from U-^13^C-glucose, which confirms their endogenous metabolic origin and allows us to propose the identities for them all (**Figure 3, Extended Data Figure 8**). Although a few FAs belong to low-abundant, but previously described series [e.g. the n12 (**Extended Data Figure 8c, g, l**) and n13 series (**Extended Data Figure 8h, m, r**)] ^27^, most correspond to well-known series, and a detailed reconstruction of their metabolic route is possible in all cases (**Figure 3b-d, Extended Data Figure 8**). For example, we detect and calculate synthesis parameters for five FA(18:2) (18:2n6, nv, nx, ny, nz). Based on their retention time and expected n-series, v, x, y and z should be > 6 (**Extended Data Figure 7i**). FA(18:2nz) is identified as FA(18:2n10) because SCDi does not affect any synthesis parameter and FADS2 decreases the calculated *E*_*1*_ (i.e. *E*_*1*_*’* = *E*_*1*_*\*Δ*_*1*_) (**Extended Data Figure 8i**). For FA(18:2nv) and FA(18:2nx), SCD1i decreases the calculated *S* more than *E*_*1*_, but the opposite occurs for FA(18:2ny). Thus FA(18:2nv, nx) and FA(18:2ny) are respectively identified as FA(18:2n7) and FA(18:2n9) (**Figure 3b-d**). Based on the FADS2i inhibition profile, we conclude that FADS2 introduces the second double bond at the 18-carbon level for FA(18:2nv) because FADS2i decreases the calculated *E*_*1*_ more than the calculated *S*, and at the 16-carbon level for FA(18:2nx) because FADS2i decreases *the* calculated *S*. Therefore, the four unknown FA(18:2) are identified as FA(18:2n7)(Δ6,11), FA(18:2n7)(Δ8,11), FA(18:2n9)(Δ6,9) and FA(18:2n10)(Δ5,8), respectively (**Figure 3b-d** and **Extended Data Figure 8i**). To facilitate the application of this strategy, we have built a decision tree that guides the identification of each double bond position based on the inhibition profile (**Extended Data Figure 9**). Eleven of the proposed identities are confirmed with commercially available standards (**Extended Data Figure 10**), and our proposed strategy discloses a more comprehensive FA biosynthetic landscape, including the description of the synthesis route of nine FAs whose proposed identities do not match previously described FAs (**Figure 4, Extended Data Figure 8**).

**Figure 3.**
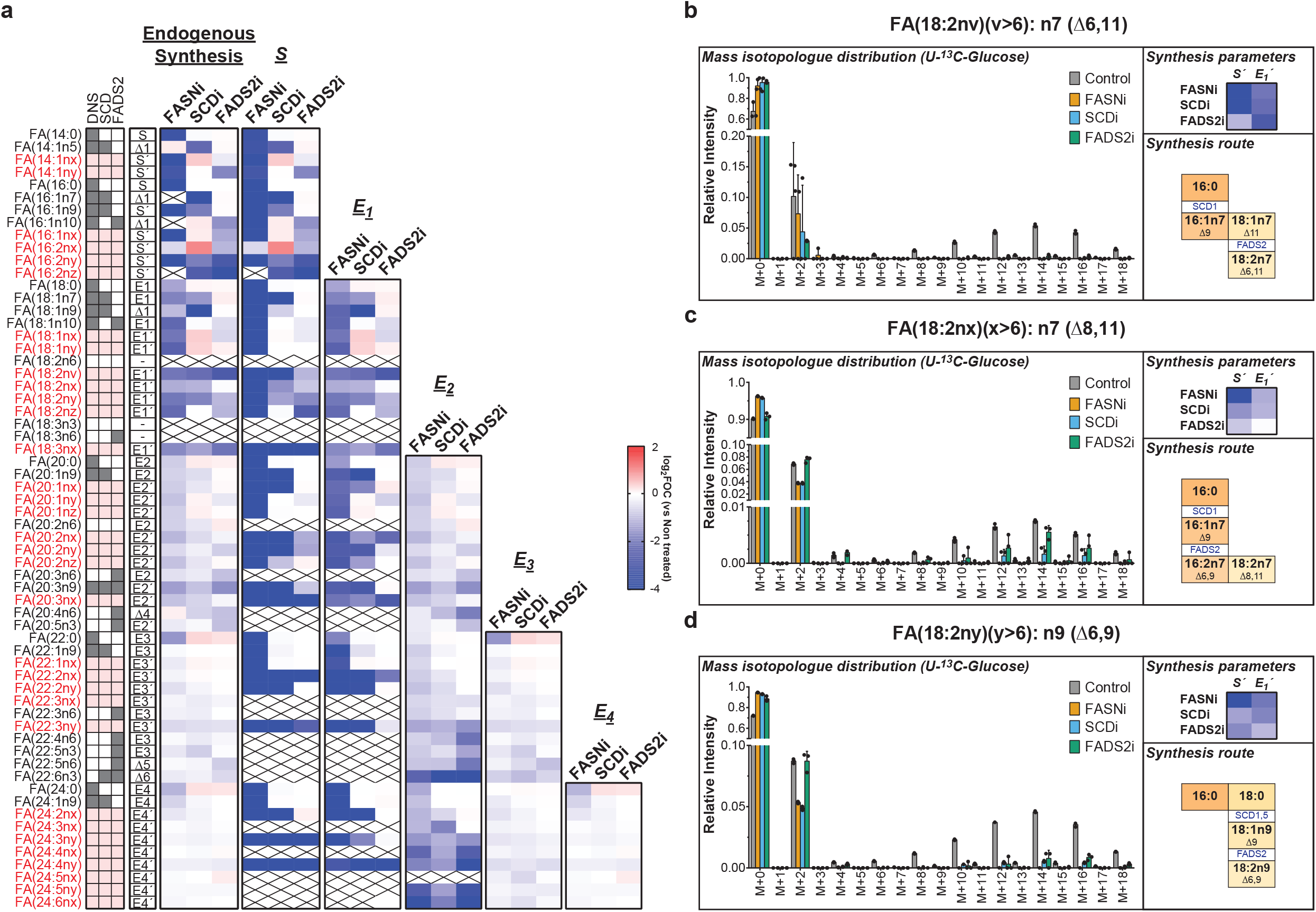
Elucidation of the synthesis route of unidentified FA species by combining FAMetA and FA metabolism inhibitors. Analysis of alterations in the FA metabolic network in the human NSCL cell line A549 incubated for 72 h with ^13^C-glucose induced by FASN inhibitor GSK2194069, SCD inhibitor A93572 and FADS2 inhibitor SC26196. **a**, Heatmap showing the mean value of the log_2_ fold-of-change (vs. untreated) for each detected FA in the following parameters: endogenously synthesised fraction, calculated *S, E*_*1*_, *E*_*2*_, *E*_*3*_ and *E*_*4*_. For each FA, the parameter reported for the endogenous synthesis is indicated. The shadowed cells indicate the activities (DNS, SCD or FADS2) involved in the synthesis of a particular FA. Red denotes the FAs whose synthesis route is unknown. On the heatmap, crosses indicate missing or NA values. **b-d**, The mass-isotopologue distribution, the mean value of the log_2_ fold-of-change (vs. untreated) in the synthesis parameters, and the proposed synthesis route for FAs FA(18:2nv) (**b**), FA(18:2nx) (**c**) and FA(18:2ny) (**d**), whose identities do not match any standard employed for the method development. In all cases n=3. Individual points are shown for the mass isotopologue distributions. The mean values are reported elsewhere.In the synthesis route description, horizontal transitions denote elongations (enzymes not indicated) and vertical transitions depict desaturations (enzymes indicated). See **Extended Data Figure 3** as a guide as to how calculations are done and which parameters are reported as the endogenous synthesis of each detected FA.

## Discussion

The therapeutic inhibition of specific FA metabolic enzymes/transporters has been proposed in diseases like cancer ^21,23,28–30^, non-alcoholic fatty liver disease ^31^, autoimmunity ^32^ or viral infection ^33^. Metabolic plasticity in FA desaturation has been recently acknowledged as a relevant phenomenon that supports lipid biosynthesis ^25,27^ and confers a metabolic advantage upon SCD inhibition in cancer cells ^25^. The expression of particular elongases (e.g. ELOVL2 in glioma ^34^ or ELOVL5 in prostate cancer ^35^) supports cell growth, tumour initiation and metastasis. Despite the wide variety of FAs, their biosynthetic routes and proven functions, current state-of-the-art tools/algorithms do not provide a comprehensive characterization of FA metabolism. The most commonly used algorithm (i.e. ISA) was initially develped for the determination of DNL for FA(14:0) and FA(16:0) ^11,12^. Further developments enabled the estimation of elongations ^13,14^ and of the de novo synthesis of odd-chain FAs ^36^. Additionally, a simple strategy for the estimation of the desaturation of FA(18:1n9) based on the ratio of the total labelling of FA(18:0) and FA(18:1n9) was discussed ^15,19^. The shown relevance of long FAs, the importance of desaturation in cell biology and the physiopathology of many diseases and the lack of a tool that performed a comprehensive characterization of all the biosynthetic reactions within FA metabolism in a user-friendly platform accessible to the broad lipid metabolism community have motivated us to develop FAMetA.

Our results demonstrate that FAMetA deciphers both patterns of global changes (**Figure 2b-c**) and detailed information about alterations in the synthesis route of FAs of interest (**Figure 2d-g**) both *in vitro* and *in vivo* (**Extended Data Figure 6**). Strikingly, the use of U-^13^C-glucose and well-characterised inhibitors of FA metabolism enzymes (i.e., FASN, SCD1, and FADS2), combined with FAMetA data analysis, enables the comprehensive characterisation of the FA biosynthetic network in A549 cells (**Figure 3, Extended Data Figure 8**). It also discloses the identity of 12 novel FAs that belong to already described n-series, which extends the known FA biosynthesis network compared to previous tools (**Figure 4**). Lack of well-characterised inhibitors of FADS1 or elongases (ELOVL1-7) limits the level of detail that can be achieved when identifying FAs by their metabolic reconstruction (**Extended Data Figure 9**). Likely some detected FAs, which are identified as the product of double desaturation introduced by the consecutive action of SCD1 and FADS2, are instead a mixture in which the products of a double desaturation introduced by SCD1 and FADS1 are also present. So the unambiguous identification of the proposed unknown/novel FAs would require using complementary analytical tools and, if possible, authentic chemical standards. Nevertheless, we demonstrate that FAMetA enables the straightforward mapping of FA biosynthetic pathways by the techniques and reagents routinely used in metabolism studies.

**Figure 4.**
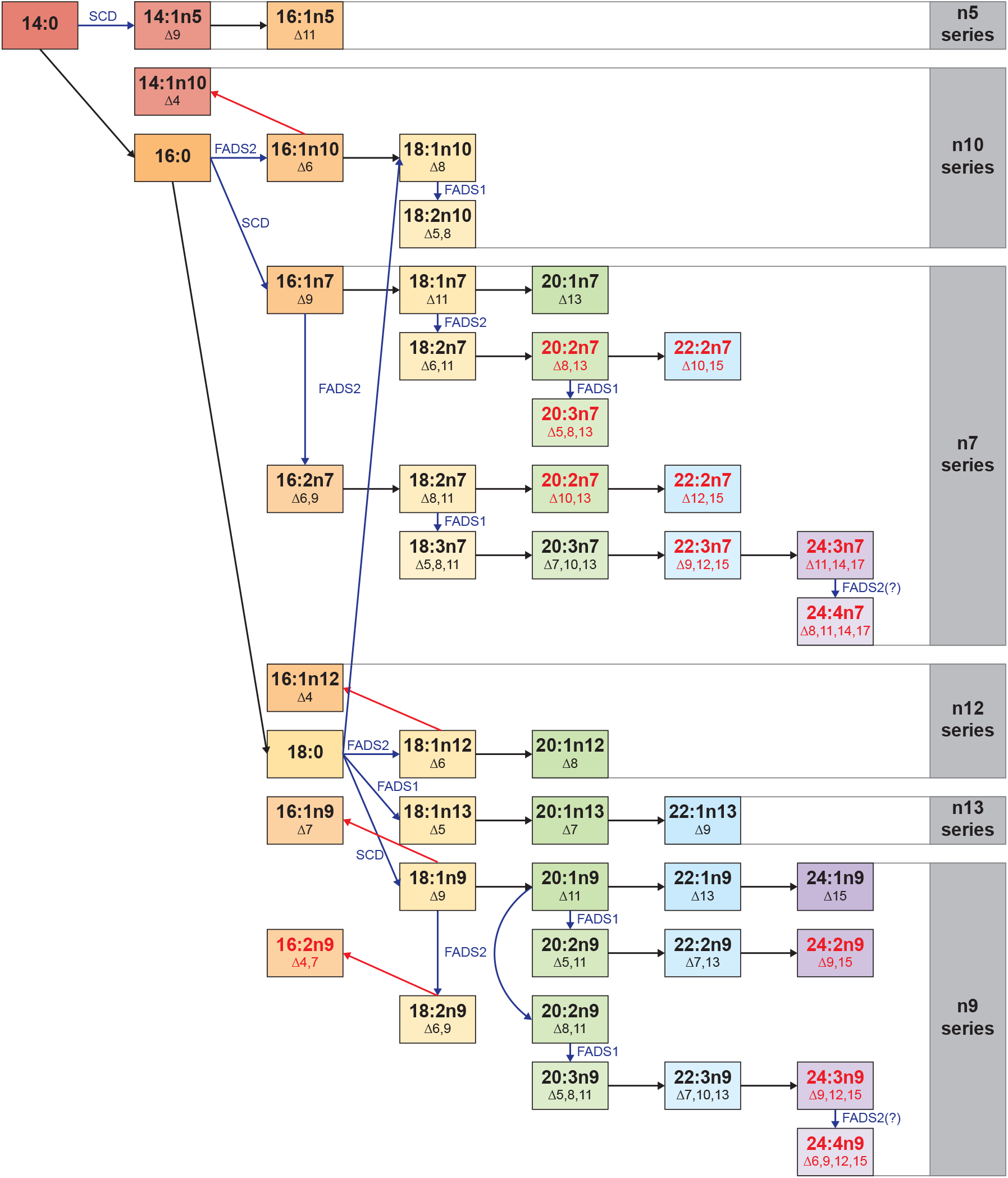
FA biosynthesis routes in the NSCLC cell line A549. Summary of the FA metabolism network in the A549 cells for those FAs that come from DNL. Black arrows denote elongations, blue arrows desaturations (the responsible enzyme is indicated) and a red arrow degradation. Red depicts the FAs that have not been previously described.

Compared to previous tools FAMetA offers (**Supplementary Table 1**): 1) the characterization of a broader FA biosynthesis network as it includes in a single tool DNL, elongation and desaturation; 2) the possibility of running all the required steps from data pre-processing to analysis of FA metabolism and graphical representation in a single tool; 3) a user-friendly environment thanks to its implementation as R-package and web-based app; 4) better fitting to the experimental data thanks to the implementation of quasi-multinomial fitting that incudes de parameter *Φ* that accounts for data overdispersion; 5) better description of elongations, thus enabling an easier interpretation of the estimated parameters; 6) easy-to-interpret parameters and graphical representations that lead to obtain meaningful biological conclusions.

Future developments of mass isotopologue data analysis tools, including FAMetA, should address some unresolved issues like using labelled-FAs as nutrients, distinguishing the uptake of exogenous FA and the lipolysis of stored lipids, estimating the synthesis rate of the FAs that result from the degradation of a longer FA [e.g. FA(16:1n9), where *S’*=*S*\**E*_*1*_\**degradation*], or the resolution of the FA metabolism properties of particular lipid classes of interest or organelles. Additionally, the FAMetA algorithm is exclusively designed to fit the data from ^13^C-based tracers for even-chain FAs. Thus future efforts should focus on implementing calculations based on ^2^H-tracers, such as ^2^H_2_O, which contributes to FA synthesis via direct H_2_O incorporation, and also via NADPH ^7,8^, and to expand the reactions to cover odd-chain FAs, in which not only the lipogenic Acetyl-CoA has to be estimated, but also the lipogenic Propionyl-CoA pool ^36^. Despite these limitations, and as far as we know, FAMetA constitutes the first tool that enables reliable estimations of FA import, synthesis, elongation and desaturation for the whole FA metabolic network of FAs within the range from 14 to 26 carbons. The FAMetA workflow includes all the required functionalities (data preprocessing, FA metabolism analysis, group-based comparisons, graphical representation) to run a complete data analysis on a single platform (**Figure 1**). Its combination with the systematic genetic manipulation of enzymes/transporters involved in FA metabolism can contribute to the characterisation of FA metabolism in unprecedented detail. Finally, to spread its use, FAMetA is freely available as an open-source R package and a web-based application (www.fameta.es). In conclusion, we believe that FAMetA is a valuable addition to existing tools and has the potential to become a key resource to study the complex FA biosynthetic landscape.

## Online Methods

### Reagents

Standard chemical reagents were obtained from Sigma-Aldrich. RPMI 1640 with stable glutamine (ref L0498) and 100x streptomycin/penicillin solution (ref L0010) were obtained from Biowest. Foetal bovine serum (FBS, ref A3160502) and dialysed FBS (ref 26400044) were obtained from Gibco. RPMI 1640 media without glucose, glutamine, and amino acids (ref R9010-01) were supplied by USBiological. U-^13^C-glucose, U-^13^C-glutamine and U-^13^C-glutamine were obtained from Cambridge Isotope Laboratories. FASN inhibitors GSK2194069 ^21^ and FADS2 inhibitor SC26196 ^24^ were purchased from Sigma-Aldrich; SCD inhibitor A93572 ^22,23^ was obtained from MedChemExpress. FA standards came from Sigma-Aldrich, Larodan and Cayman Chemicals. Antibodies anti-CD3 (ref BE0001-1) and anti-CD28 (ref BE0015- 1) were provided by BioXCell. Recombinant IL-2 (ref 212-12) was obtained from Peprotech.

### Mice

For all the experiments, 8-10-week-old female mice were used. 6-week-old wild-type C57BL/6 were purchased from Charles River Laboratories. Mice were left in a normal light cycle (08:00–20:00h) and had free access to water and a standard chow diet. Animals were housed in the Health Research Institute– Hospital La Fe Valencia facilities. Mouse studies followed the protocols approved by the Health Research Institute–Hospital La Fe Valencia Ethics and Animal Care and Use Committee (Protocol number 2020/VSC/PEA/0048).

### Isolation, culture, and stimulation of mouse naïve CD8+ T-cells

To isolate naïve CD8^+^ T-cells, spleens were harvested. Single-cell suspensions were prepared by manual disruption and passage through a 70- μm cell strainer in PBS supplemented with 0.5% BSA and 2 mM EDTA. After RBC lysis, naïve CD8+ T-cells were purified by magnetic bead separation using commercially available kits following manufacturers’ instructions (naïve CD8a+ T-Cell Isolation Kit, mouse, Miltenyi Biotec Inc.)^37^.

Cells were cultured in complete RPMI media (RPMI 1640 supplemented with 10% FBS, 100 U ml−1 penicillin, 100 μg ml−1 streptomycin, 55 μM 2-mercaptoethanol). Naïve T-cells were stimulated for 48 h with plate-bound anti-CD3 (10 μg ml−1) and anti-CD28 (5 μg ml−1) in complete RPMI media supplemented with recombinant IL-2 (100 U ml−1). All the experiments on ‘active’ T-cells were performed on day 4–5 postactivation^37^.

### Cell lines and growth conditions

The KRAS-mutant NSCLC cell line A549 was originally obtained from ATCC. The A549 cells were maintained in RPMI-1640 media supplemented with 10% FBS, 100 U ml−1 penicillin and 100 μg ml−1 streptomycin, and were routinely screened for mycoplasma contamination. Identity was confirmed by STR sequencing.

### Cell metabolism studies

sotopically-labelled media were prepared from glucose, glutamine and amino acids-free RPMI media, and were supplemented with 10% dialysed FBS, 100 U/ml penicillin and 100 μg/ml streptomycin. For the culture of the CD8^+^ T-cells, media were also supplemented with recombinant IL-2 (100 U ml−1) and 55 μM 2-mercaptoethanol. U-^13^C-glucose and U-^13^C-glutamine were added at the normal concentration found in RPMI 1640 media. U-^13^C-acetate was added at 100 μM. U-^13^C-lactate was added at 11 mM ^37–41^.

The CD8^+^ T-cells were seeded at 0.8 × 10^6^ cells/mL and incubated for 72h with labelled media (and inhibitors). At 24h and 48h, cells were counted using the Countess II automated cell counter (Thermo Fischer Scientific) and density was adjusted to 0.8 × 10^6^ cells/mL with complete fresh labelled media (and inhibitors). At 72h, the final cell density was determined. Then,cells were transferred to 1.5 mL Eppendorf tubes and pelleted (500 g, 3 min). Media were removed. Cells were washed once with cold PBS 1x, resuspended in 500 μL of cold PBS 1x and stored at -80°C ^37,39^.

For the NSCLC cell line A549, cells were seeded at 7 × 10^4^ cells/well in 6-well plates. After 24 h, media were replaced with labelled media (and inhibitors). Cells were incubated for 48-72 h until 80-90% confluence, the media was replaced with fresh media (and inhibitors) every 24h. At the end of the incubation, media were removed, cells were washed once with cold PBS 1x, scraped with 500 μL of cold PBS 1x, transferred to 1.5 mL Eppendorf tubes and stored at -80°C ^38^.

### Saponification and extraction of total Fas

To analyse the total FAs, 450 μL of cell suspension were transferred to a glass vial, and 1,000 μL of a 9:1 MeOH:KOH (3M in H2O) solution containing PC(16:0/16:0)D62 at 3 ppm were added. Saponification was performed for 1 h at 80° in a water bath. After saponification, samples were cooled on ice and acidified by adding 100 μL of formic acid. FAs were extracted with 2 mL of heptane:isooctane (1:1) (2x), dried in a nitrogen flow, resuspended in 200 μL of mobile phase A containing myristic acid D27 at 1 ppm and transferred to a glass HPLC vial ^42^.

### UPLC-HRMS analysis of FAs

FAs were analysed in a quadrupole–orbitrap mass spectrometer (Q Exactive, Thermo-Fisher Scientific) coupled to reverse phase chromatography via electrospray ionisation. Liquid chromatography separation was performed in a Cortecs C18 column (2.1 mm × 150 mm, 1.6 μm particle size; Waters). Solvent A was 2.5 mM ammonium acetate in 60:40 water:methanol. Solvent B was 2.5 mM ammonium acetate in 95:5 acetonitrile:isopropanol. The flow rate was 300 μL/min, the column temperature was 45°C, the autosampler temperature was 5°C and the injection volume was 5 μL. The liquid chromatography gradient was: 0 min, 45% B; 0.5 min, 45% B; 19 min, 55% B; 23 min, 99% B; 34 min, 99% B. Between injections, the column was washed for 2 min with 50:50 acetonitrile:isopropanol before being equilibrated to the initial conditions. The mass spectrometer operated in the negative-ion mode to scan from m/z 100 to 450 at a resolving power of 140000. Data were acquired in the centroid mode.

### FAMetA

#### FAMetA implementation

FAMetA was developed in an R programming environment. It is available *via* CRAN (https://CRAN.R-project.org/package=FAMetA). In addition, the web-based implementation of FAMetA was built using the Shiny R-package (Shiny: Web Application Framework for R. 2021). It is accessible at www.fameta.es (**Supplementary Figure 11**).

#### The FAMetA workflow

The FAMetA workflow starts with raw MS data files in the mzXML format, which can be obtained with any MS file converter, e.g. msConvert from ProteoWizard ^43^, and a csv file containing the required metadata (sample name, acquisition mode, sample group, or class, and any additional information like external measures for normalisation) (**Supplementary Figure 2, steps 1-2**). Data preprocessing can be performed in the R environment/web-based application using our proposed workflow, which combines functions from FAMetA and our previously described R-package LipidMS ^44,45^ (available *via* CRAN (https://CRAN.R-project.org/package=LipidMS)) (**Supplementary Figure 2, steps 2-5**).

LipidMS is called for the first preprocessing step, which runs peak-peaking, alignment and grouping through functions *batchdataProcessing, alignmsbatch* and *groupmsbatch* (**Supplementary Figure 2, step 2**). Then FAMetA is called, and functions *annotateFA* and *curateFAannotations* are used to identify any unique FA isomers. Automatic FA annotations can be exported to a csv file and be modified by removing rows of unwanted FA by modifying the initial and end retention times, or by adding new rows with missing compounds. Unique compound names with nomenclature “FA(16:1)n7”, where n7 (omega-7) indicates the last double-bond position, are required to differentiate FA isomers. For any unknown positions, letters x, y and z are allowed (i.e. FA(16:1)nx). The internal standards for later normalisation can also be added in a new row at this point by indicating IS in the compound name column (**Supplementary Figure 2 step 3**). Once FAs have been correctly identified, FA isotopes can be extracted using function *searchFAisotopes* (**Supplementary Figure 2, step 4**). Finally, data can be corrected and normalised using the *dataCorrection* function, which runs four different steps (all of which are optional): data correction for natural ^13^C abundance using the accucor algorithm^46^; data normalisation with internal standards; blank subtraction; external normalisation (**Supplementary Figure 2, step 5**). Alternatively, the external data processed by other available software/tools can be loaded at this workflow point or before the data correction and normalisation steps.

Then the actual FA metabolism analysis can be performed by sequentially running the *synthesisAnalysis, elongationAnalysis* and *desaturationAnalysis* functions (**Supplementary Figure 3, steps 1-3**). The first two functions model isotopologue distributions by non-linear regression (https://CRAN.R-project.org/package=minpack.lm) with many initial values ^47^ to ensure that the best fits are found. By default, a maximum of 1,000 iterations for synthesis and 10,000 for elongation are performed for each set of initial values to fit the isotopologue distributions (maxiter parameter) or until the model has converged 100 times (maxconvergence parameter). If no results are obtained or parameters come close to the limits of the confidence intervals, these parameters can be increased to improve the results. The third function employs the previous results to estimate the desaturation values. Finally, the summarised results tables and heatmaps are obtained using the *summarizeResults* function to export and explore the results (**Supplementary Figure 3, step 4**).

#### Estimation of the DNL parameters

We considered FA(16:0) the final DNL product. Thus FAMetA can estimate the DNL parameters for FAs up to 16 carbons. For these species, *I* and *S* represent the fraction of the FA pool that is synthesised and imported, respectively, and sums 1:

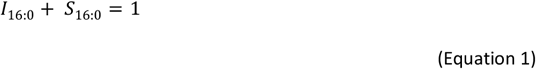

For the DNL analysis, FA isotopologue distributions (previously corrected for the natural abundance of the ^13^C isotopes) are modelled with the following sum of the weighted quasi-multinomial distributions adapted from ^48^:

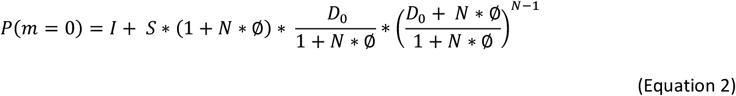

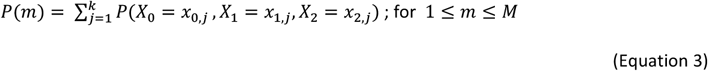

where:

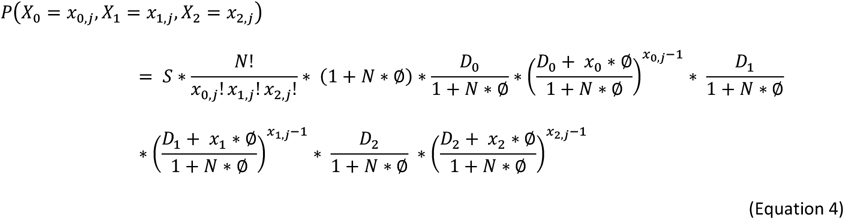

given that:

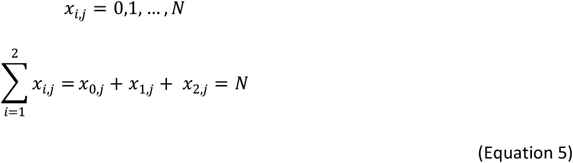

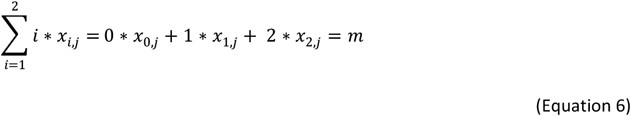

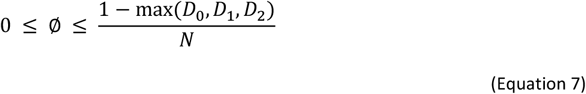

*M* is the total number of carbons in the FA molecule and *N* = *M*/2. This represents the number of acetyl-CoA molecules used for the synthesis of an FA of length M. m is the number of ^13^C atoms incorporated into the FA molecule. *D*_*0*_, *D*_*1*_ and *D*_*2*_ represent the fraction of acetyl-CoA with 0, 1 or 2 atoms of ^13^C, respectively, and sum 1. *x*_*0*_, *x*_*1*_ and *x*_*2*_ represent the number of acetyl-CoA units with 0, 1 or 2 ^13^C atoms that provide an *M*-carbon FA with an *m* label. For a given pair of *N* and *m* values, up to *k* combinations of the *x*_*0*_, *x*_*1*_ and *x*_*2*_ values fulfil Equations 5 and 6. *Φ* accounts for overdispersion and can be set at 0 to reduce quasi-multinomial distribution to multinomial distribution. The *in silico* validation of the above-described equations demonstrates an overestimation of *Φ* and an underestimation of *S* and *D*_*2*_ for values of D_2_ ≥ 0.75. In these situations, the upper limit of *Φ* is set at 0.5*(1-max(D_0_, D_1_, D_2_)/N. Note that overdispersion parameter *Φ* modifies *D*_*0*_, *D*_*1*_ and *D*_*2*_ for each synthesis step, which allows distribution to widen or narrow. Based on this model, non-linear regression (https://CRAN.R-project.org/package=minpack.lm) with many sets of plausible initial values (adapted from ref ^47^) is used to fit the observed isotopologue distributions of FAs up to 16 carbons, and to estimate parameters *D*_*1*_, *D*_*2*_, *Φ* and *S*. To improve the analysis results, distributions FA(16:0) and FA(14:0) are firstly fitted, and the estimated parameters D_1_, D_2_ and Φ are used to model the other FAs.

#### Elongation

The main product of the DNS of FA is FA(16:0) ^1^. So the main DNS route, plus elongation, starts at 16 carbons and then adds blocks of two carbons. Elongation from FA(14:0) is a minor route^14^ and is omitted for simplicity. For the FAs ranging from 18 to 26 carbons, the following equations are considered:

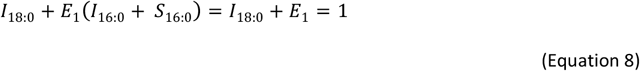

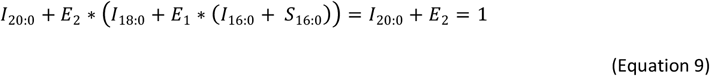

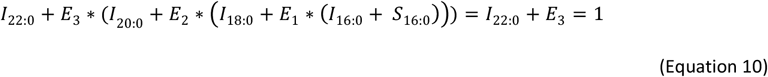

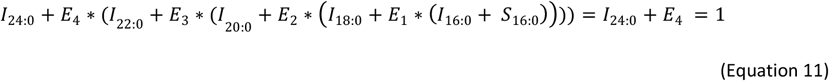

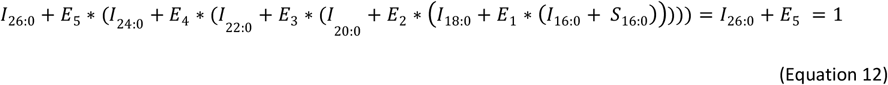

For the elongation analysis of endogenous FA, isotopologue distributions are modeled using Equation 2 for synthesis until FA(16:0), followed by single independent elongation steps (*E*_*1*_, *E*_*2*_ …, *E*_*n*_). The probability of incorporating 0, 1 or 2 ^13^C atoms into the FA to be elongated equals *E*_*i*_*D*_*0*_, *E*_*i*_*D*_*1*,_ and *E*_*i*_*D*_*2*_, respectively. Parameters *D*_*0*_, *D*_*1*_ and *D*_*2*_ are imported from FA(16:0)/FA(14:0). Hence the only relevant parameters to be estimated in the elongation analysis are *E*_*i*_ and *I*. For FA(18:0), FA isotopologue distributions (previously corrected for natural ^13^C isotopes abundance) are modelled with the following equations:

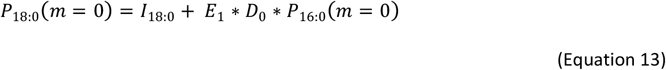

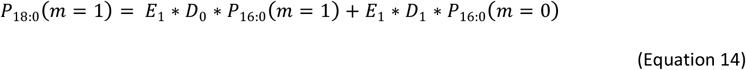

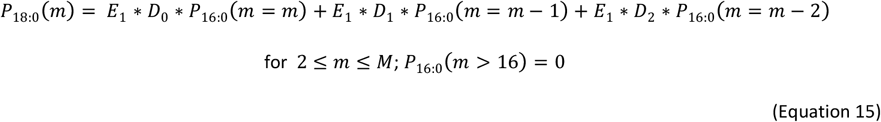

Analogous equations can be obtained for FA with M>18 by adding elongation terms to previously existing distributions. For series n6 and n3 (**Extended Data Figure 1**), elongation is usually expected from FA(18:2)n6 and FA(18:3)n3. Thus synthesis (*S*_*16:0*_) and the first elongation step (*E*_*1*_) are set at 0. If isotopologue M+2 is observed given the degradation of FA(18:2)n6 or FA(18:3)n3, followed by one elongation step, then *E*_*1*_ is estimated. However, the endogenously synthesised fraction remains at NA. Once again, non-linear regression (https://CRAN.R-project.org/package=minpack.lm) with multiple initial values ^47^ is used to fit the observed isotopologue distributions of the elongated FAs.

#### Desaturation

After estimating the synthesis and elongation parameters, these results can be used to calculate the FA fraction that comes from desaturation in the unsaturated FA. For a given unsaturated FA (e.g. FA(18:1n9)), we can conceptually consider a one-step elongation-desaturation reaction (in this example, directly from FA(16:0) to FA(18:1n9)), or a two-step elongation followed by a desaturation process (in this example FA(16:0) is elongated to FA(18:0) and then desaturated to FA(18:1n9)):

**(Scheme 1).**
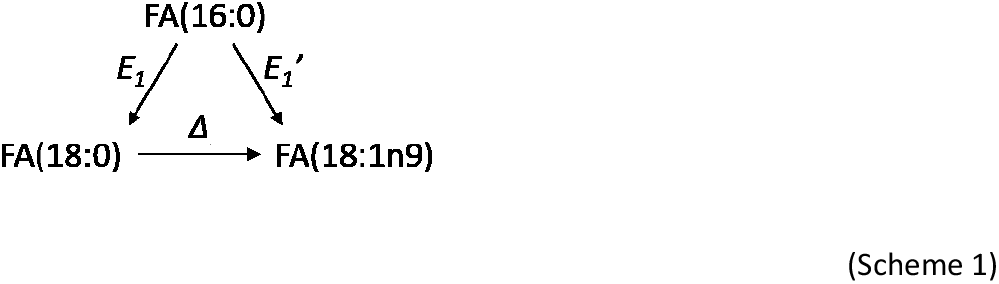

By using FAMetA, we can directly estimate both *E*_*1*_ and *E*_*1*_*’* from the isotopologue distributions of FA(18:0) and FA(18:1n9), respectively. From alternative paths, the relative import and endogenous synthesis pathways of FA(18:1n9) can be written as:

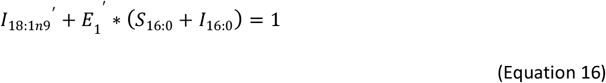

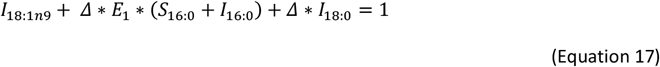

By combining both equations, we can define that:

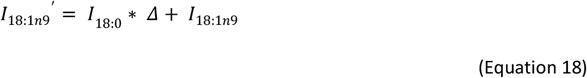

and, thus, calculate desaturation parameter *Δ* as:

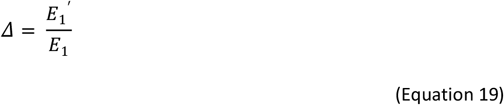

If both *E*_*i*_*’* and *E*_*i*_ are below the confidence interval, which is set at 0.05 by default for desaturation, parameter *Δ* is not calculated, and *E*_*i*_*’* remains as the endogenously synthesised fraction. If the stationary state is not reached, values > 1 can be obtained for the desaturation parameter that is, in this case, replaced with 1.

This same model can be used for all the known desaturation steps provided that the precursor and product FA isomers are correctly and uniquely identified, and the stationary state is reached. For the FA synthesised from desaturation activities, Δ is considered the fraction from endogenous synthesis. So the imported fraction is calculated as 1-Δ. With unknown isomers or missing precursors, S’ or E’ is returned for the *DNS* of FAs until 16 carbons or the elongation of longer FAs, respectively. The reactions included in FAMetA are described in **Extended Data Figure 1** ^14,16,49,50^. However, additional reactions (desaturations) can be included for unknown/additional FAs by modifying *desaturationdb* in FAMetA.

#### In silico tests of FAMetA

To test FAMetA’s performance with different FA isotopologue distributions and noise levels, *in silico* tests on models are run. To evaluate FAMetA’s performance to estimate parameters for the DNS analysis, realistic values for *D*_*1*_ (5 values from 0 to 0.2), *D*_*2*_ (15 values from 0 to 1), *Φ* (10 values from 0 to 0.1) and *S* (15 values from 0 to 1) are combined to simulate 3,945 theoretical FA(16:0) distributions to which 0%, 2%, 5% and 10% noise levels are added to obtain 10 different noised distributions for each set of parameters. Bias (evaluated as an absolute or relative error) and dispersion (evaluated as RSD) are calculated and graphically represented for parameters *D*_*2*_, *S* and *Φ* (**Supplementary Figure 4**).

To evaluate FAMetA’s performance to estimate the parameters for the elongation analysis, the mass isotopologue distributions for FA(18:0), FA(20:0), FA(22:0), and FA(24:0) are generated. To evaluate the elongation of FA(16:0) to FA(18:0), *D*_*1*_ and *Φ* are set at 0.05 and 0.01, respectively. The realistic values for *D*_*2*_ (9 values between 0.1 and 0.9), *S* (19 values between 0.05 and 1) and *E*_*1*_ (19 values between 0.05 and 1) are employed to generate 3,249 theoretical FA(18:0) distributions. For FA(20:0), FA(22:0) and FA(24:0), the synthesis parameters for FA(16:0) are set at D_1_ = 0.05, Φ = 0.01 and S = 0.6. Nine values within the 0.1-0.9 range and 10 values within the 0.1-1 range are generated for *D*_*2*_ and *E*_*n*_, respectively. Bias (evaluated as a relative error) and dispersion (evaluated as RSD) are calculated and graphically represented for all the estimated parameters (**Supplementary Figures 5-8**).

To evaluate FAMetA’s performance to estimate the parameters for the desaturation analysis, the mass isotopologue distributions for FA(16:1n7), and FA(18:1n9) are generated. *D*_*1*_ and *Φ* are set at 0.05 and 0.01, respectively. For FA(16:1n7), 13 values within the 0.1-0.87 range and 14 values within the 0.07-1 range are generated for *D*_*2*_ and *S*, respectively. For FA(18:1n8), 13 values within the 0.1-0.87 range and 14 values within the 0.07-1 range are generated for *D*_*2*_ and *E*_*1*_, respectively. In both cases, 14 values within the 0.07-1 range are generated for Δ. Bias (evaluated as a relative error) and dispersion (evaluated as RSD) are calculated and graphically represented for parameter *Δ* for both FAs (**Supplementary Figure 9**).

## Data availability

The input mzXML LC-MS data files used to generate **Figures 2-3**, the metadata associated with the studies, the FA identities for each dataset, the preprocessed data ready to be used to estimate the FA metabolism parameters for each study and an R script are available at Zenodo with accession number 6511248.

## Code availability

FAMetA’s source code is offered to the public as a freely accessible software package under the GNU GPL license, version 3. It is available at https://github.com/maialba3/FAMetA and Zenodo with accession number 6511248.

## Acknowledgments

We thank Prof. Cholsoon Jang (University of California, Irvine) for kindly providing the raw data to generate Extended Data Figure 6 and for his input to the manuscript. M.A.A.-B. is supported by a PFIS contract from the Carlos III Health Institute of the Spanish Ministry of Economy and Competitiveness (FI18/00224). J.C.G.-C. is supported by a grant from the Conselleria de Sanidad Universal y Salud Pública, Generalitat Valenciana, as part of Plan GenT, Generació Talent (DEI-01/20-C). A.L. is supported by the European Regional Development Fund (FEDER) and the Carlos III Health Institute of the Spanish Ministry of Economy and Competitiveness (PI20/00580 and DTS2019/0143). Part of the equipment used in this work was co-funded by the Generalitat Valenciana and European Regional Development Fund (FEDER) funds (PO FEDER of Comunitat Valenciana 2014-2020).

## Author Contributions

J.C.G.-C and A.L. conceived the method. A.L. supervised study development. M.I.A.-B. designed and programmed the algorithms for FAMetA and its online implementation. M.I.A.-B. and J.C.G.-C. optimised the LC-MS-based method for the FA analysis. M.I.A.-B. and J.C.G.-C. performed the FA analyses. J.C.G.-C. and M.B. performed the labelling experiments. M.I.A.-B., J.C.G.-C., M.B., O.J., and A.L. analysed the data and discussed the results. J.C.G.-C. and A.L. wrote the manuscript in collaboration with all the authors.

## Competing Interests statement

O.J. reports receiving honoraria for advisory roles from Boehringer Ingelheim, Bristol-Myers Squibb, Merck Sharp & Dohme, Roche/Genetech, AstraZeneca, Pfizer, Eli Lilly, AbbVie. A.L. reports receiving honoraria for advisory roles from AstraZeneca.

**Extended Data Figure 1.**
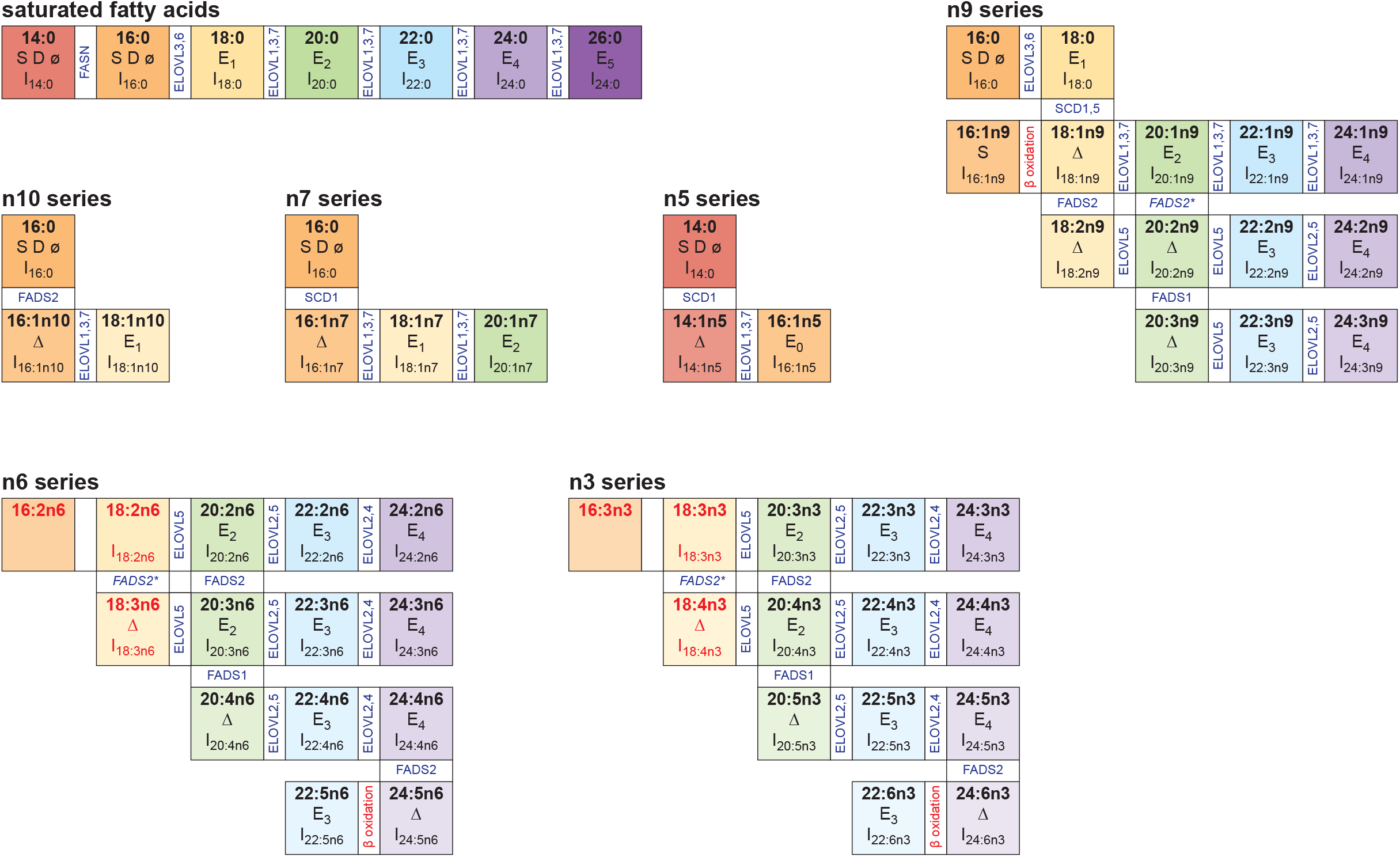
FA metabolism network. Summary of the FA interconversions covered by FAMetA and the parameters that can be estimated for each one. In red, the FA for which no parameter can be estimated because they are either solely imported or result from desaturation being performed on them. Horizontal transitions denote elongations and vertical transitions depict desaturations. The responsible enzymes are indicated in both cases. We assume DNL up to FA(16:0), although the calculation of the DNL parameters can be estimated for both FA(14:0) and FA(16:0). For the transformations of FA(18:2n6) into FA(20:3n6) and FA(18:3n3) to FA(20:4n3), the preferred route is desaturation, followed by elongation. The asterisk denotes a secondary route.

**Extended Data Figure 2.**
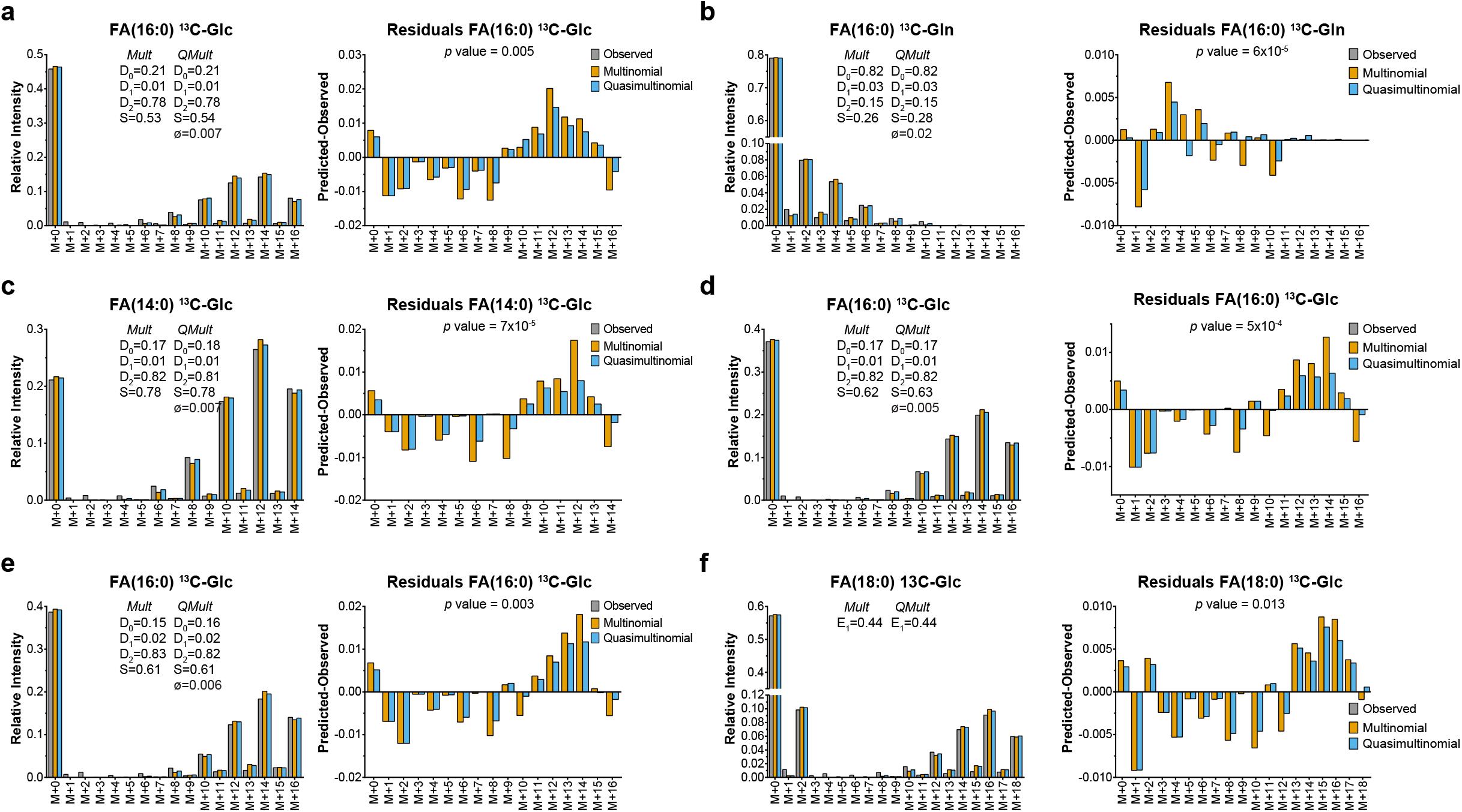
Fitting experimental mass-isotopologue FA data to multinomial and quasi-multinomial distributions. **a-b**, FA(16:0) in the A549 cells upon incubation with U-^13^C-glucose (**a**) or U-^13^C- glutamine (**b**), data obtained from ref.^18^. **c-d**, FA(14:0) (**c**) and FA(16:0) (**d**) in the H1299 cells upon incubation with U-^13^C-glucose, data obtained from ref.^14^. **e-f**, FA(16:0) (**e**) and FA(18:0) (**f**) in the MCF7 cells upon incubation with U-^13^C-glucose, data obtained from ref. ^13^. For each dataset, the experimental data, the fitting done using the FAMetA algorithm with multinomial or quasi-multinomial distributions, and the residuals are shown. The reported *p* values correspond to the comparisons between multinomial and quasi-multinomial fitting using a log-likelihood ratio test and right-tailed chi-square distribution.

**Extended Data Figure 3.**
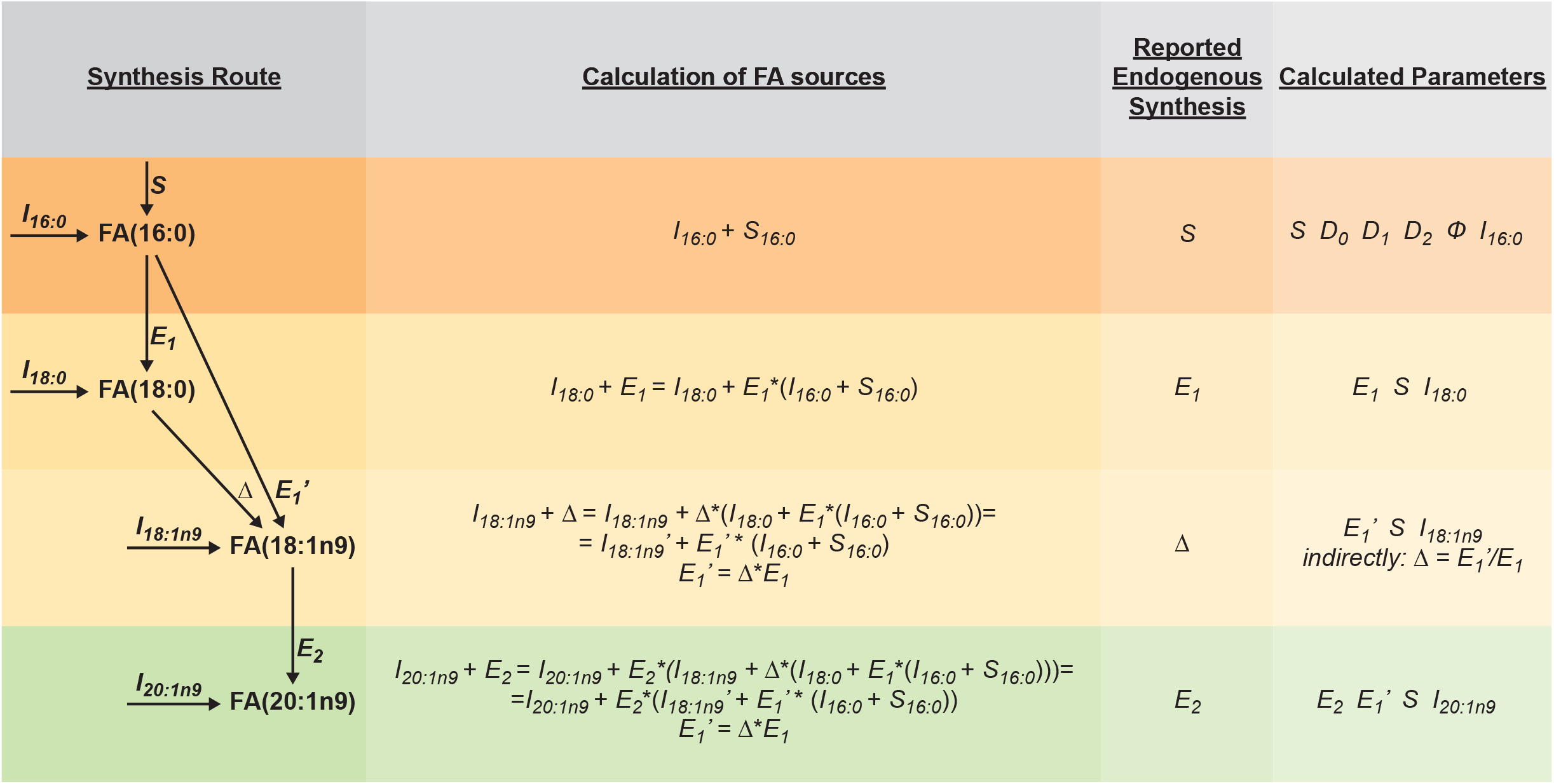
Example of the FAMetA calculations for FA(16:0) to FA(20:1n9). A detailed description of the calculation of FA sources, reported endogenous synthesis and the parameters calculated for the FAs FA(16:0), FA(18:0), FA(18:1n9) and FA(20:1n9).

**Extended Data Figure 4.**
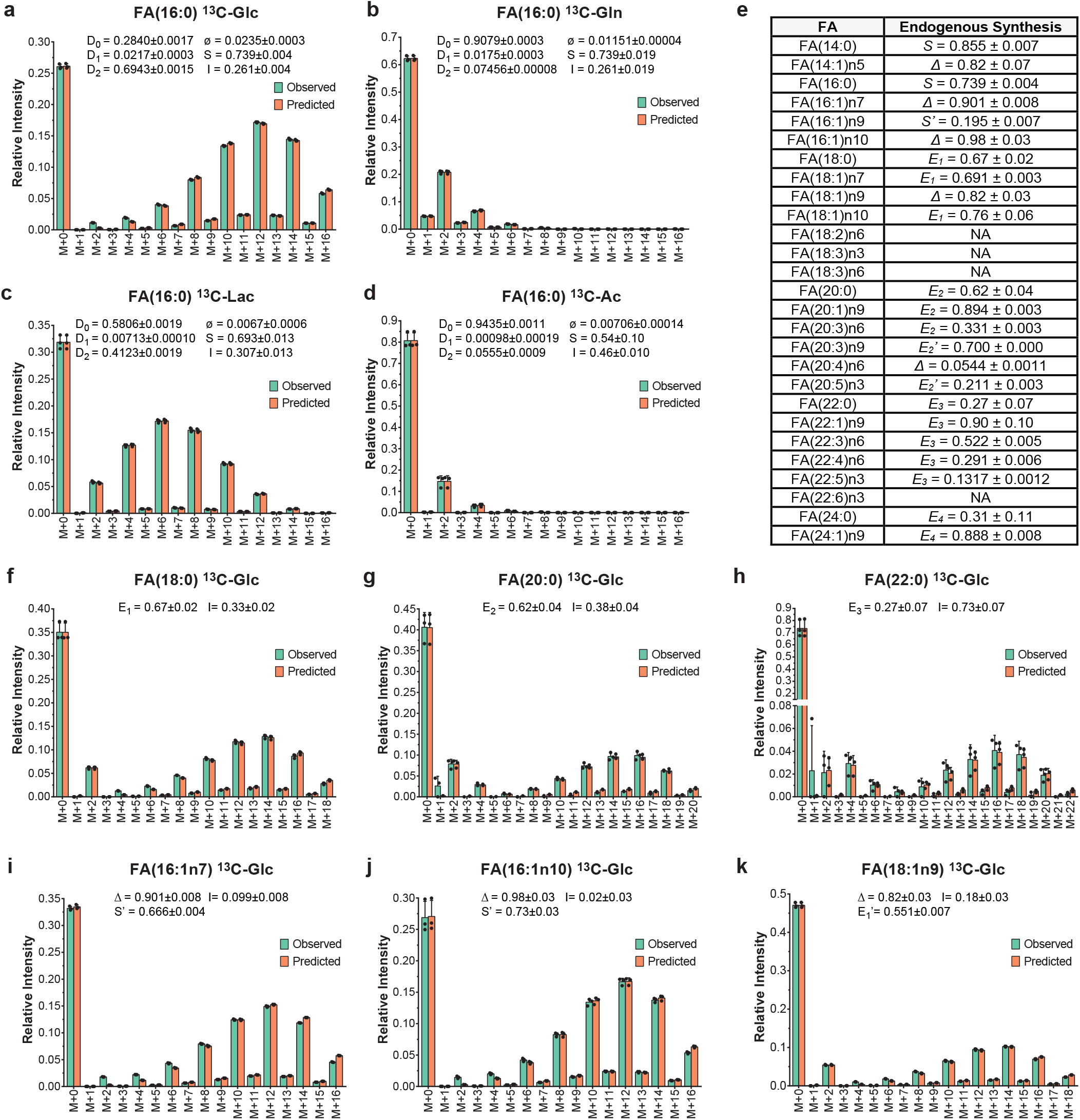
Biological validation of FAMetA in active mouse CD8^+^ T-cells. Estimation of the FA metabolism parameters in the active mouse CD8^+^ T-cells incubated for 72 h with various U-^13^C-tracers. **a-c** Estimation of the sources and the DNL parameters for FA(16:0) upon incubation with U-^13^C-glucose (**a**), U-^13^C-glutamine (**b**), U-^13^C-lactate (**c**) or U-^13^C-acetate (**d**). **e-k** Estimation of the synthesis parameters for various FAs upon incubation with U-^13^C-glucose. **e** Summary of the endogenously synthesised fraction for the 27 known FAs detected in the active mouse CD8^+^ T-cells. **f-k** Estimation of the sources for FA(18:0) (**f**), FA(20:0) (**g**), FA(22:0) (**h**), FA(16:1n7) (**i**), FA(16:1n10) (**j**), FA(18:1n9) (**k**). Data are expressed as mean±SD (n=3).

**Extended Data Figure 5.**
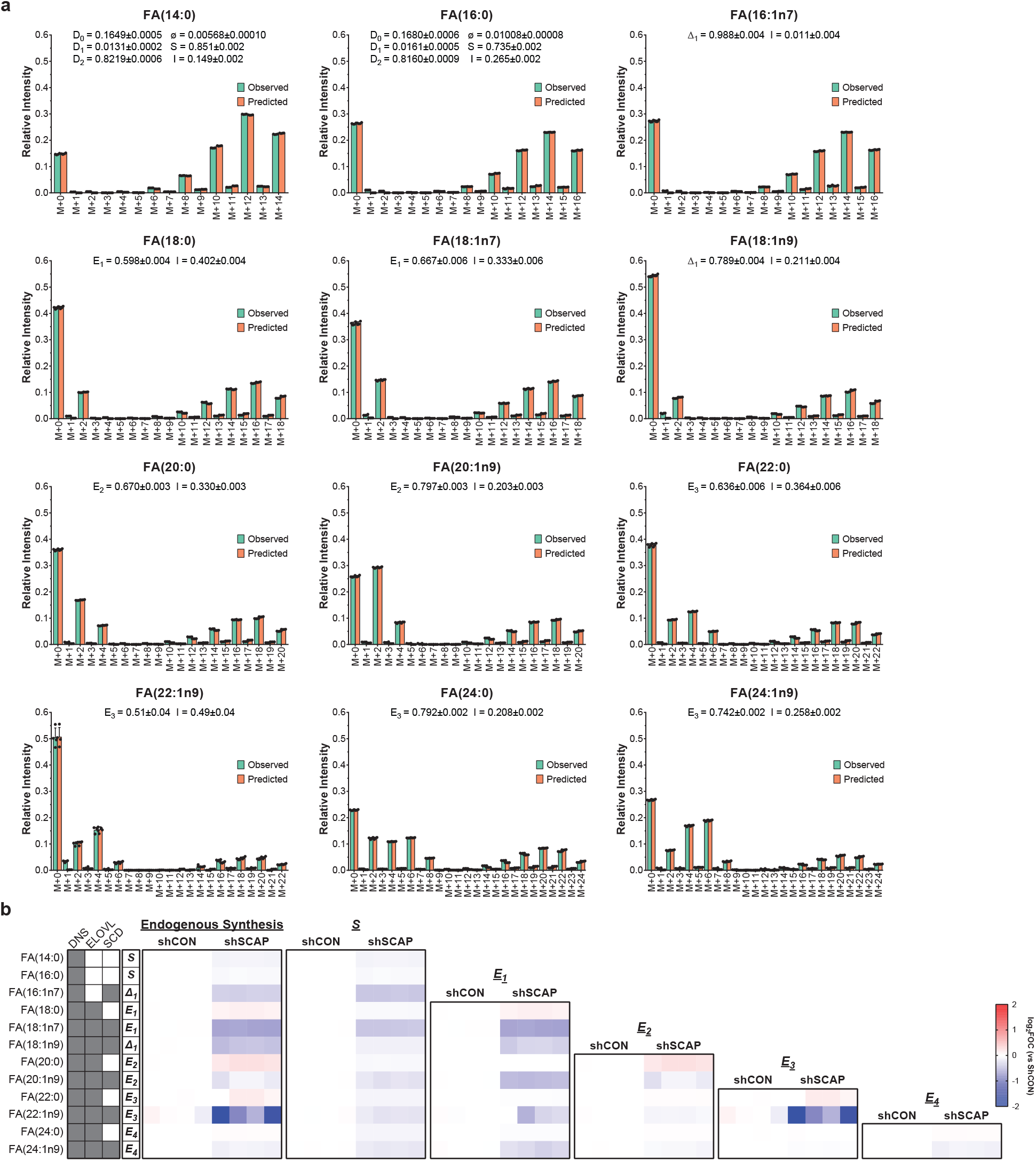
Analysis of the influence of the down-regulation of SCAP on the FA metabolism in the H1299 cells; data obtained from ref.^14^. **a**, FAMetA was used to fit all the reported experimental mass-isotopologue distributions for the control condition (shCON). **b**, Heatmap showing the log_2_ fold of change (vs. shCON) for each reported FA in the following parameters: endogenously synthesised fraction, calculated *S, E*_*1*_, *E*_*2*_, *E*_*3*_ and *E*_*4*_. For each FA, the parameter reported for endogenous synthesis is indicated. The shadowed cells indicate the activities (DNS, elongation (ELOVL), or SCD1-mediated desaturation (SCD)) involved in the synthesis of a particular FA. See **Extended Data Figure 3** as a guide as to how calculations are done and which parameters are reported as the endogenous synthesis of each detected FA. n=4.

**Extended Data Figure 6.**
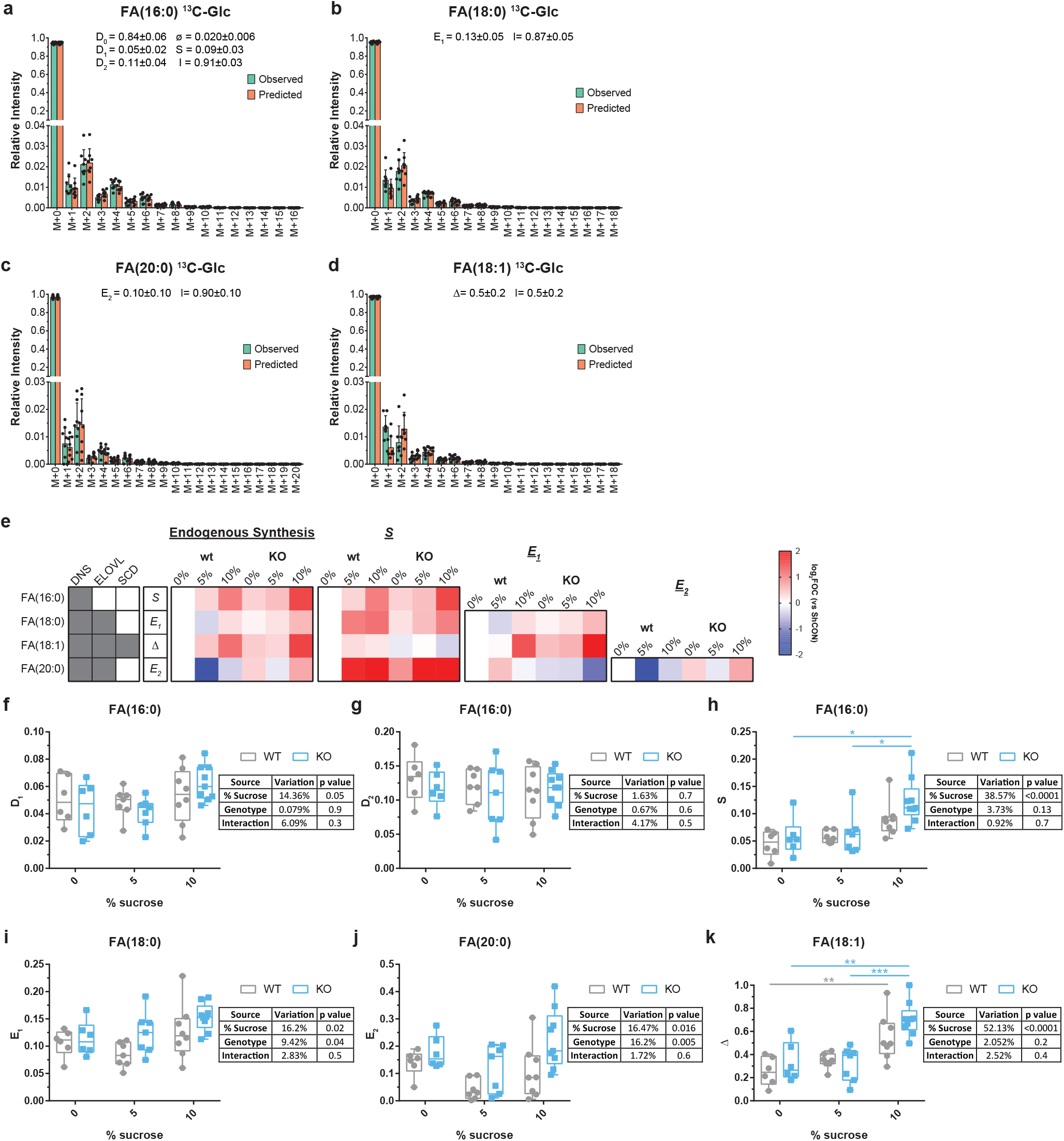
Effect of KHK-C expression and dietary sucrose on FA synthesis *in vivo*, data obtained from ref.^26^. **a-d**, FAMetA was used to fit all the reported experimental mass-isotopologue distributions for the WT 10% sucrose group. **e-j**, The calculated FA synthesis parameters obtained with FAMetA. The tables summarise the result of the two-way ANOVA performed for each calculated parameter. Paired differences are calculated by a *post hoc* Tukey test. The p-values obtained for the reported significant differences: S parameter for FA(16:0), KO 10% vs. KO 0%, p-value =0.014, KO 10% vs. KO 5%, p-value =0.03; Δ parameter for FA(18:1), WT 10% vs. WT 0%, p-value =0.006, KO 10% vs .KO 0%, p-value =0.0012, KO 10% vs. KO 5%, p-value =0.0004 (n=6,6,7,7,8,8). **k**, Heatmap showing the log_2_ fold of change (vs. WT 0% sucrose) for each reported FA in the following parameters: endogenously synthesised fraction, calculated *S, E*_*1*_ and *E*_*2*_. For each FA, the parameter reported for the endogenous synthesis is indicated. The shadowed cells indicate the activities [DNS, elongation (ELOVL) or SCD1-mediated desaturation (SCD)] involved in the synthesis of a particular FA. See **Extended Data Figure 3** as a guide as to how calculations are done and which parameters are reported as the endogenous synthesis of each detected FA.

**Extended Data Figure 7.**
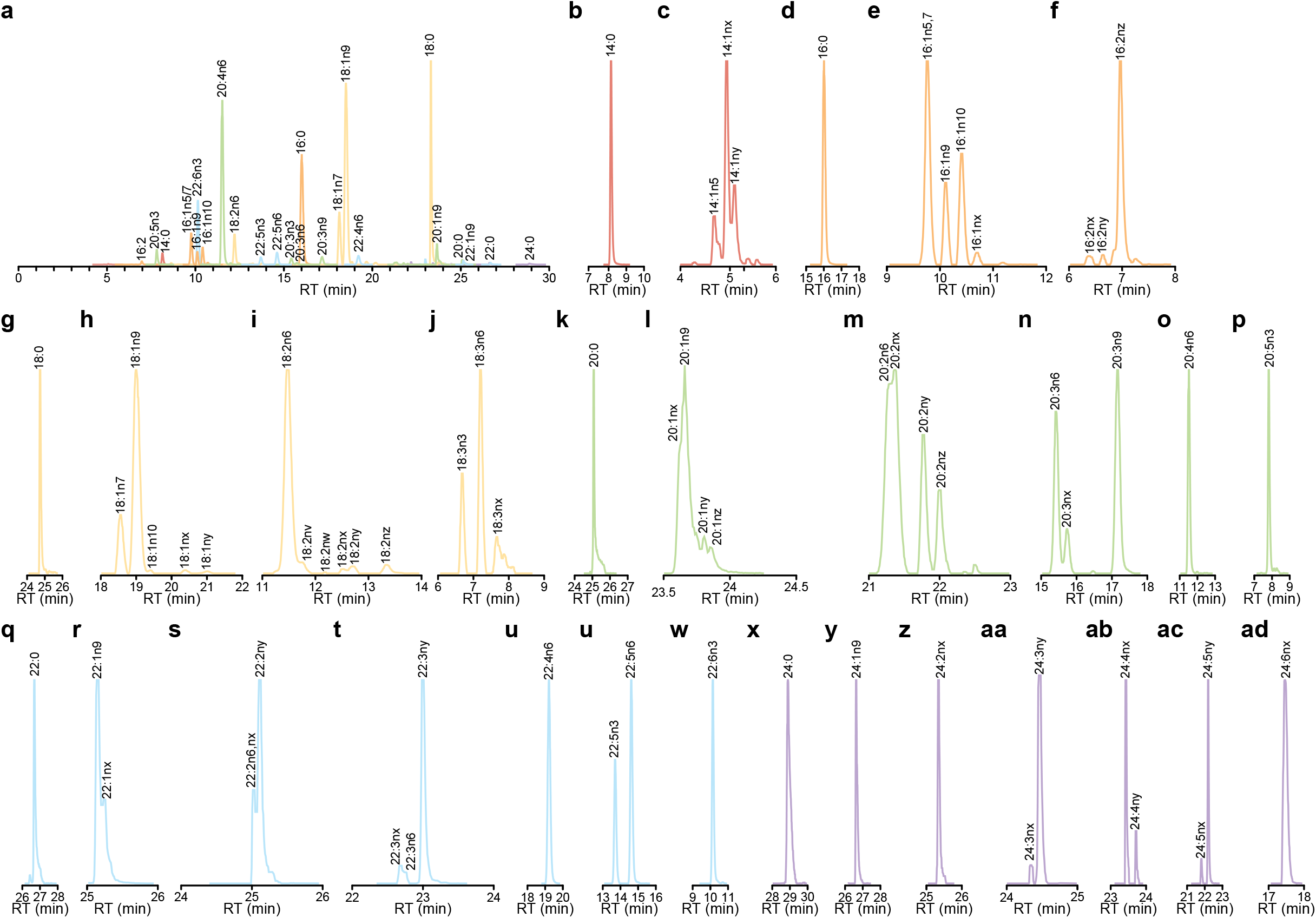
Chromatographic separation of the saponified FAs from the A549 cells in culture. **a**, all the detected FAs. **b**, FA(14:0). **c**, FA(14:1n5, nx, and ny). **d**, FA(16:0). **e**, FA(16:1n5,n7,n9, n10, nx, and ny). **f**, FA(16:2nx, ny, and nz). **g**, FA(18:0). **h**, FA(18:1n7, n9, n10, nx, and ny). **i**, FA(18:2n6, nw, nx, ny, and nz). **j**, FA(18:3n3, n6, and nx). **k**, FA(20:0). **l**, FA(20:1n9, nx, and ny). **m**, FA(20:2n6, nx, ny, and nz). **n**, FA(20:3n6, n9, and nx). **o**, FA(20:4n3, and n6). **p**, FA(20:5n3). **q**, FA(22:0). **r**, FA(22:1n9, and nx). **s**, FA(22:2n6, nx, and ny), **t**, FA(22:3nx, n6, and ny). **u**, FA(22:4n6). **v**, FA(22:5n3, and n6). **w**, FA(22:6n3). **x**, FA(24:0). **y**, FA(24:1n9). **z**, FA(24:2nx). **aa**, FA(24:3nx, and ny). **ab**, FA(24:4nx, and ny). **ac**, FA(24:5nx, and ny). **ad**, FA(24:6nx). Where FA(14:1nx, and ny), FA(16:1nx, and ny), FA(16:2nx, ny, and nz), FA(18:1nx, and ny), FA(18:2nw, nx, ny, and nz), FA(18:3nx), FA(20:1nx, ny and nz), FA(20:2nx, ny, and nz), FA(20:3nx), FA(22:1nx), FA(22:2nx, and ny), FA(22:3ny), FA(24:2nx), FA(24:3ny) and FA(24:4ny) are unknown FAs that incorporate the labelling compatible with an endogenous DNS origin, where FA(22:3nx), FA(24:3nx), FA(24:4nx), FA(24:5nx, and ny) and FA(24:6nx) are unknown FAs that incorporate labelling compatible with an endogenous origin that results from the elongation of an imported FA.

**Extended Data Figure 8.**
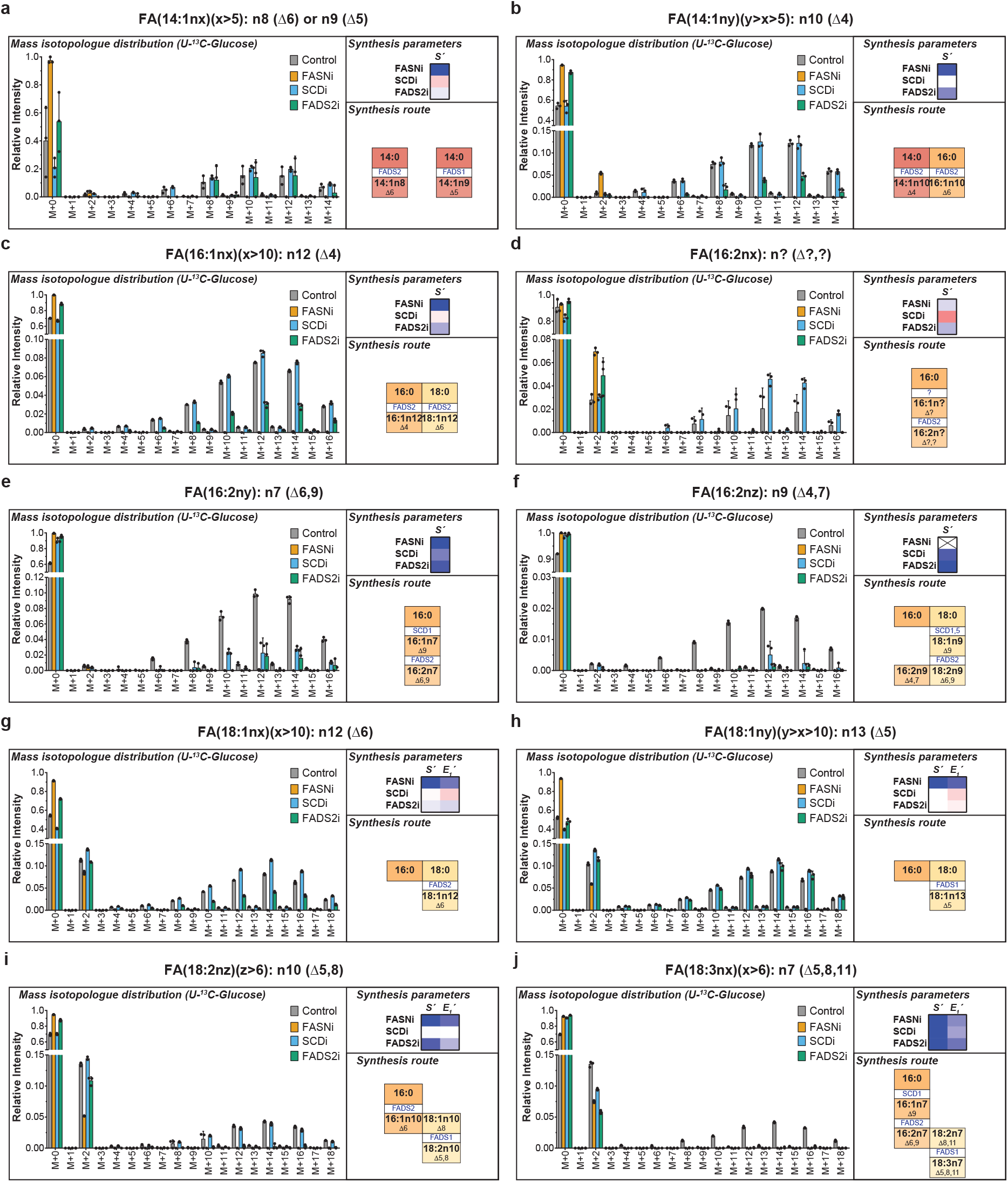
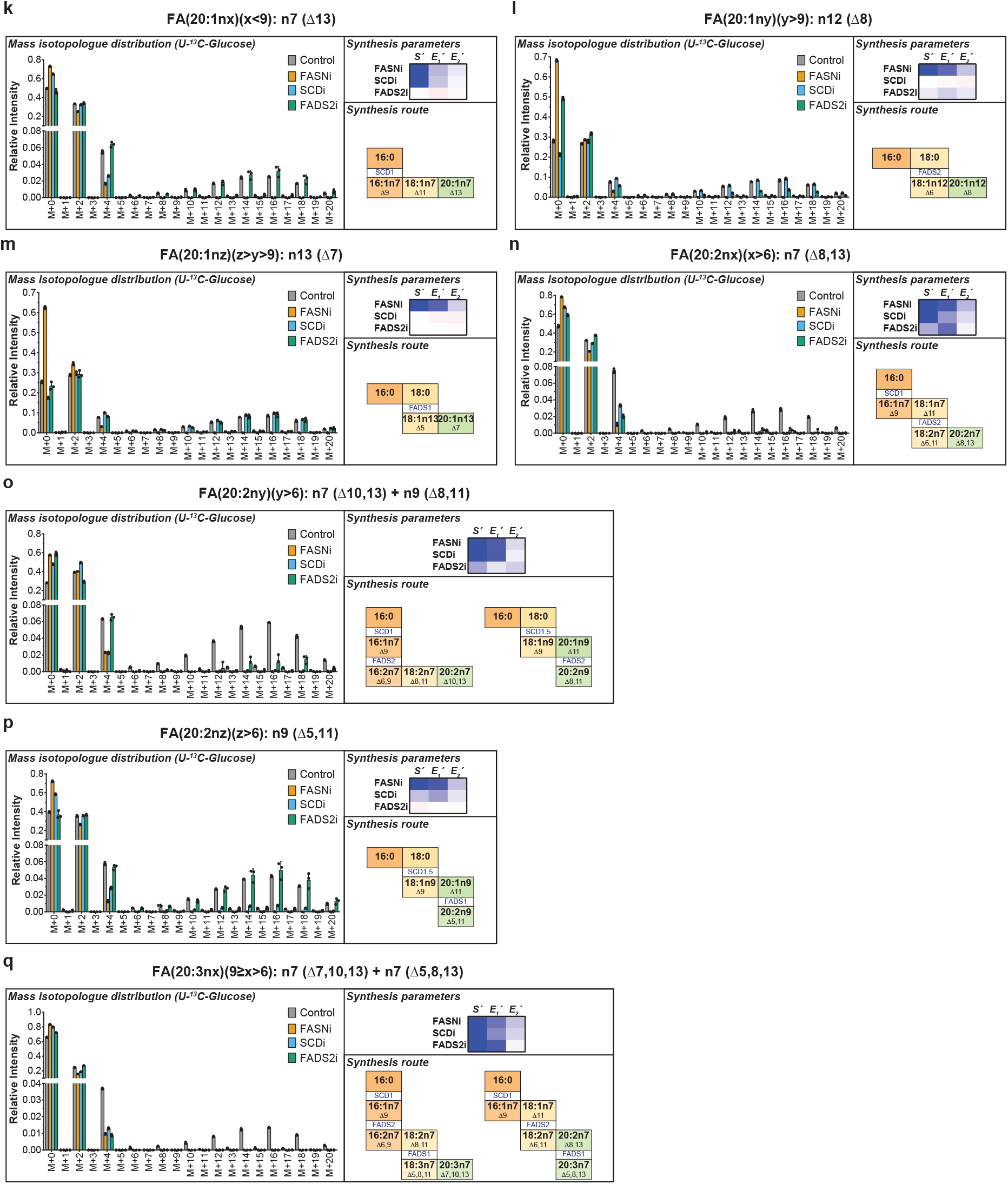
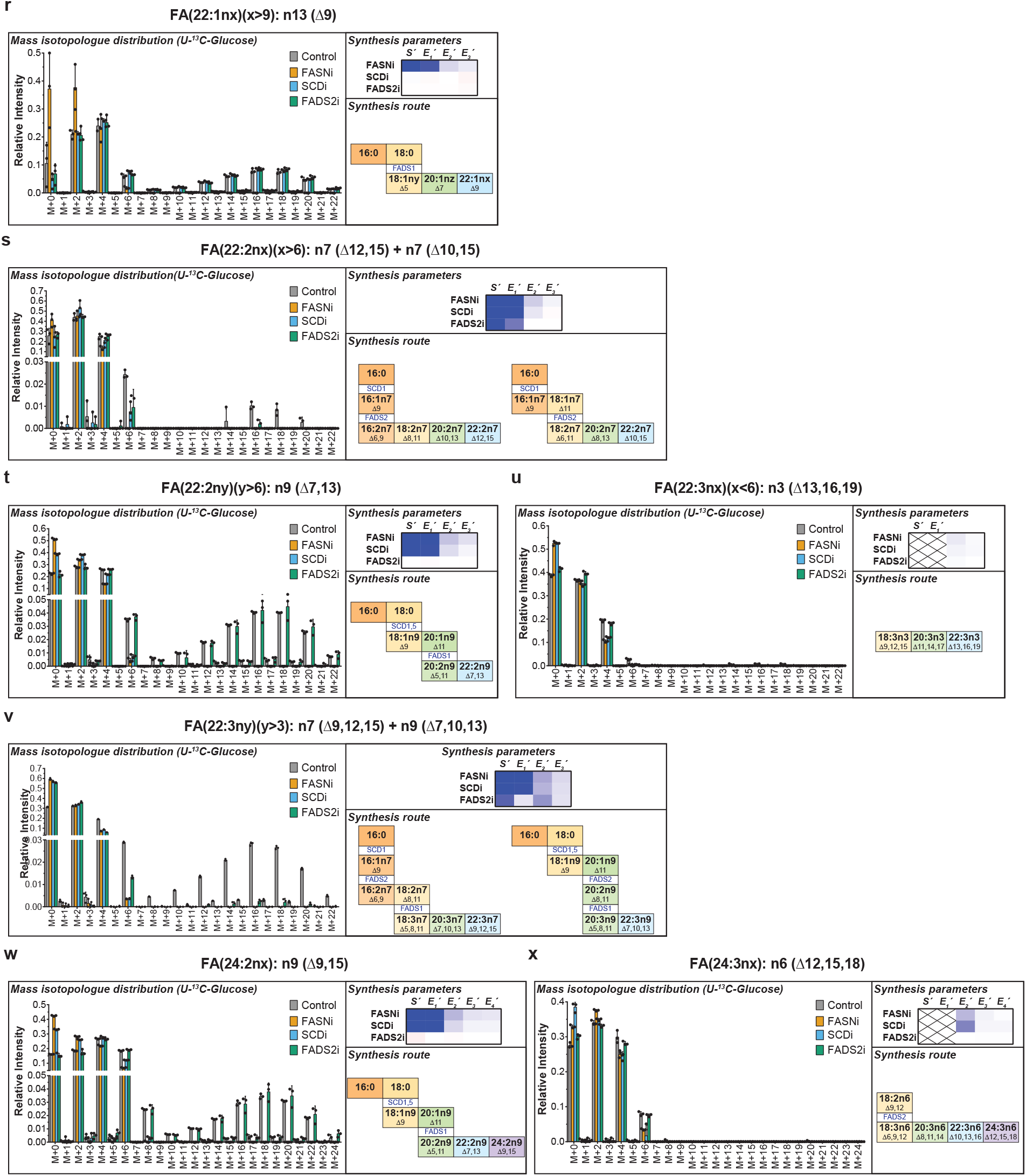
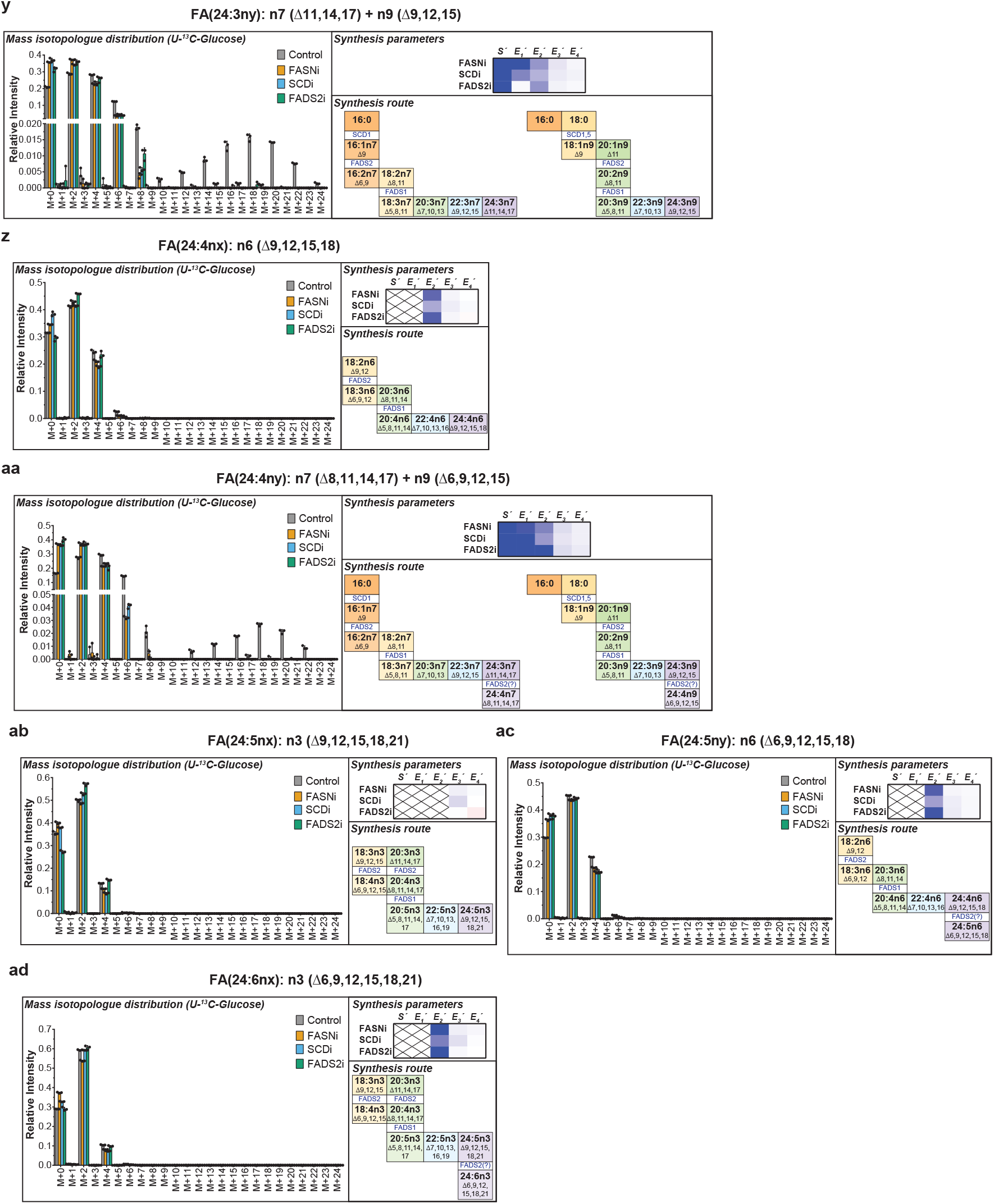
Proposed synthesis route for the unknown FAs in Figure 4. Mass-isotopologue distribution, mean value of the log_2_ fold-of-change (vs. untreated) in the synthesis parameters and the proposed synthesis route for FAs FA(14:1nx) (**a**), FA(14:1ny) (**b**), FA(16:1nx) (**c**), FA(16:2nx) (**d**), FA(16:2ny) (**e**), FA(16:2nz) (**f**), FA(18:1nx) (**g**), FA(18:1ny) (**h**), FA(18:2nz) (**i**), FA(18:3nx) (**j**), FA(20:1nx) (**k**), FA(20:1ny) (**l**), FA(20:1nz) (**m**), FA(20:2nx) (**n**), FA(20:2ny) (**o**), FA(20:2nz) (**p**), FA(20:3nx) (**q**), FA(22:1nx) (**r**), FA(22:2nx) (**s**), FA(22:2ny) (**t**), FA(22:3nx) (**u**), FA(22:3ny) (**v**), FA(24:2nx) (**w**), FA(24:3nx) (**x**), FA(24:3ny) (**y**), FA(24:4nx) (**z**), FA(24:4ny) (**aa**), FA(24:5nx) (**ab**), FA(24:5ny) (**ac**) and FA(24:6nx) (**ad**). On heatmaps, crosses indicate missing or NA values. In the synthesis route description, horizontal transitions denote elongations (enzymes not indicated). Vertical transitions depict desaturations (enzymes indicated).

**Extended Data Figure 9.**
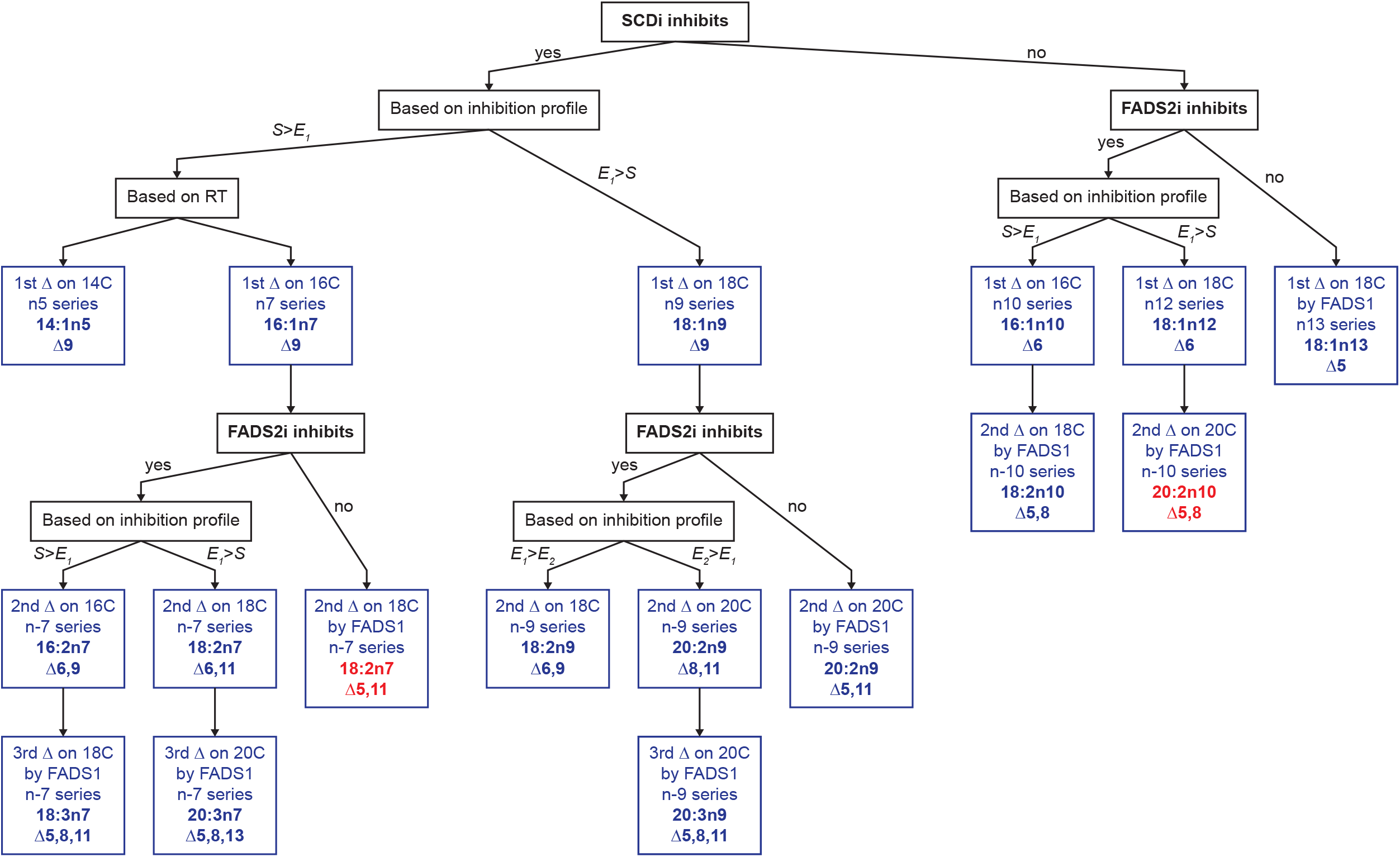
The algorithm employed to identify unknown FAs by the reconstruction of their biosynthesis route. This algorithm was used in **Figure 3** and **Extended Data Figure 8**. The depicted algorithm is applied to identify the double bond positions for FAs based on the inhibition profile obtained upon incubation with U-^13^C-glucose, either with or without SCDi or FADS2i. The algorithm applies to FAs whose origin can be tracked to FA(14:0)/FA(16:0). The previous assumptions must be met: 1) the FA incorporates labelling and intensity suffices to obtain values for all/most expected isotopomers; 2) FASNi decreases parameter S or distribution is consistent with the origin being FA(14:0)/FA(16:0). Based on the chromatographic profile, we expect the FAs to elute by increasing n-series [i.e. RT(n5 series) ≤ RT(n7 series) ≤ RT(n9 series), etc.]. The algorithm allows to identify the initial FA for the FA synthesis routes described in **Figure 4;** thus the actual position of the double bonds has to be extrapolated for the FAs of a different carbon length to that indicated in the algorithm. In red, FAs for which we can anticipate the identification and synthesis route based on the described strategy, but were not detected or unambiguously assigned experimentally in the A549 cells because FADS1 inhibitors were lacking.

**Extended Data Figure 10.**
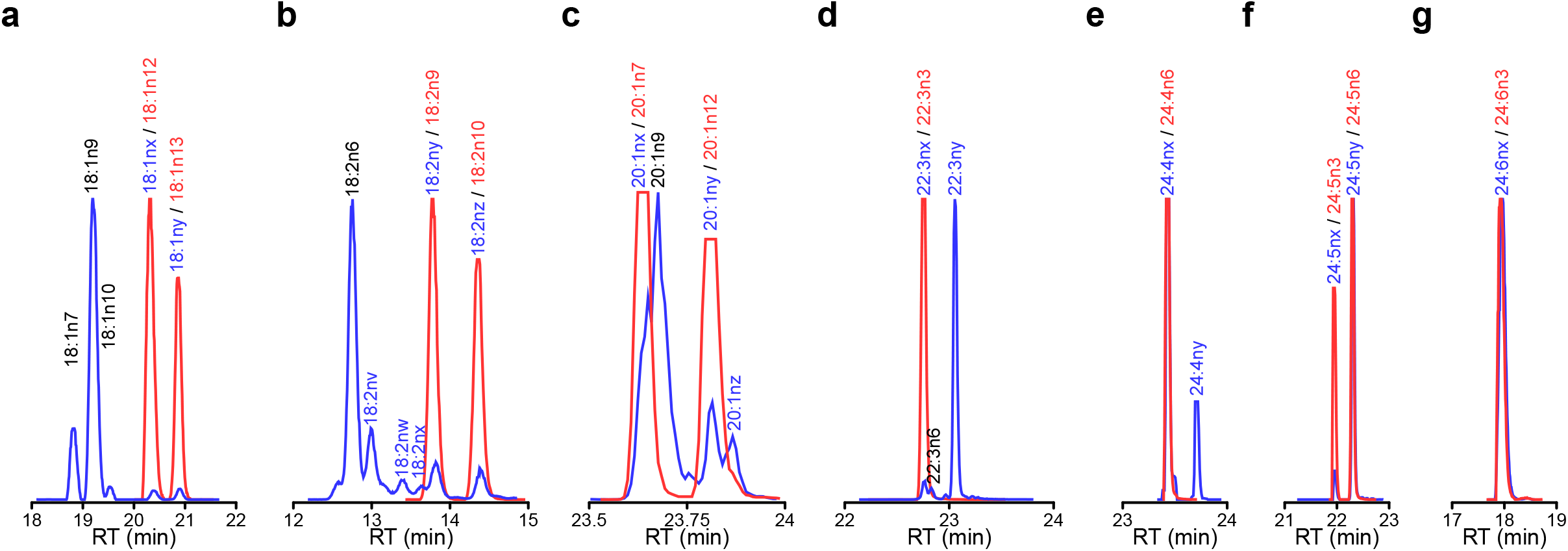
Confirmation of the identity of 11 unknown FAs in the A549 cells with chemical standards. Chromatographic separation of FAs 18:1 (**a**), 18:2 (**b**), 20:1 (**c**), 22:3 (**d**), 24:4 (**e**), 24:5 (**f**) and 24:6 (**g**). In blue, the saponified FAs from the A549 cells in culture. In red, chemical standards. Text in black, the FAs that initially matched the chemical standards used to develop the method; in blue, a notation of the unknown FAs detected in the A549 cells; in red, the chemical standards used to confirm the identity of the selected unknown FAs.

