## Supplementary Information for "FAMetA: a mass isotopologue-based tool for the comprehensive analysis of fatty acid metabolism"

#### Supplementary results

**The FA biosynthesis network in the active mouse CD8+ T-cells**. Under standard culture conditions (i.e., RPMI media and normoxia), glucose is the preferred carbon source for FA synthesis (D ≈ 0.7) in the active mouse CD8+ T-cells (**Extended Data Figure 4a**), with a minor contribution of glutamine (D ≈ 0.08) (**Extended Data Figure 4b**). If present in media, lactate and glucose are metabolically exchangeable at the lactate dehydrogenase (LDH) level. Thus, lactate feeds the pyruvate pool and FA synthesis (**Extended Data Figure 4c**). When supplemented in media, acetate feeds the acetyl-CoA pool and contributes to FA synthesis (**Extended Data Figure 4d**). These results are consistent with previously published data obtained by different approaches[18,39,41]. Twenty-seven known FAs are detected in the samples, including a variety of saturated, monounsaturated and polyunsaturated FAs within the range from 14 to 24 carbons. Although most of the identified FAs appear in culture media, endogenous synthesis is the preferential route for the saturated and monounsaturated FAs, whereas polyunsaturated FAs preferentially come from exogenous sources (**Extended Data Figure 4e-k**).

**Comparison between FAMetA and FASA.** The comparison between FAMetA and FASA is performed using a dataset published by the authors of FASA consisting of eight samples and twelve FAs in the H1229 cells incubated with U-$^{13}$C-glucose and U-$^{13}$C-glutamine either with or without the induced down-regulation of SCAP (shControl or shSCAP)[14].

We firstly compare them in computing speed terms. The processing time with Intel Xeon E5-1620 CPU (3.5 GHz) with 32 GB RAM in Windows is ~170 min for FASA (Matlab R2022a) and ~ 12 min for the FAMetA R package (R v4.1.1, RStudio v1.4, FAMetA v0.1.3). The same analysis on the FAMetA webserver takes ~30 minutes. The FAMetA algorithm calculates the fractional contribution of the carbon source ($D_0$, $D_1$ and $D_2$) and overdispersion parameter ($\Phi$) based on the distribution of FA(16:0). These values are then employed to fit the remaining FAs. The same strategy is employed for FASA by firstly fitting FA(16:0) and then the remaining FAs by setting the $D_0$, $D_1$ and $D_2$ values.

Then we compare the results obtained by both applications. FAMetA and FASA present differences in the way they calculate the FA biosynthesis parameters. While FAMetA calculates import, DNL, elongation and desaturation, FASA does not calculate desaturation. FAMetA and FASA calculate elongation by very different approaches. FAMetA provides the direct estimation of each step in a specific FA synthesis pathway (**Extended Data Figure 3**). For example, the FA(20:0) sources are described as $I_{20:0} + E_2 = I_{20:0} + E_2 * (I_{18:0} + E_1 * (I_{16:0} + S_{16:0}))$, where each parameter ($S$, $E_1$, $E_2$) directly represents a single synthesis route step, and $E_2$ is the direct estimation of the fraction of FA(20:0) that results from the elongation of the total FA(18:0) pool. Conversely in FASA FA(20:0), sources are described as $S + IE_2 + IE_1 + I$, where $S$ (elongated from FA(16:0)) actually represents $S_{16:0} * E_1 * E_2$; $IE_2$ (elongated from the imported FA(16:0)), $I_{16:0} * E_1 * E_2$; $IE_1$ (elongated from FA(18:0)), $I_{18:0} * E_2$, and $I$ represents the fraction of the directly imported FA(20:0) [14]. Using FASA, the authors of the original study conclude that SCAP down-regulation decreases both DNL and elongation[14]. Although they do not report the detailed results of each synthesis parameter for every reported FA, we analyse the dataset using FASA to find that several parameters change for each FA, and it is difficult to ascertain clear patterns to provide a more detailed conclusion than that proposed by the authors. However, thanks to the way elongations are calculated in FAMetA, the calculation of desaturations, and the proposed graphical representations in the FAMetA workflow, we identify that the main change induced by the down-regulation of SCAP is reduced SCD activity (**Extended Data Figure 5**).

Thus we conclude that compared to FASA, FAMetA provides a more comprehensive characterisation of the FA biosynthetic network, a better and more intuitive description of each synthesis parameter, and a more complete workflow that goes from data preprocessing to group-based comparisons and graphical representation. It is also more efficient from a computing perspective.

#### Supplementary Tables

**Supplementary Table 1. Comparison of the features implemented within the main available tools for the analysis of FA metabolism.**

| | **ISA**[11,12] | **ConvISA**[13] | **Kamphorst**[15,19,42] | **FASA**[14] | **FAMetA** |
|---|---|---|---|---|---|
| De novo lipogenesis | ✓ | ✓ | ✓ | ✓ | ✓ |
| Contribution of labeled nutrient to lipogenic AcetylCoA pool | ✓ | ✓ | ✓ | ✓ | ✓ |
| Elongation | ✗ | ✓ (orange) | ✓ | ✓ | ✓ |
| Desaturation | ✗ | ✗ | ✓ (orange) | ✗ | ✓ |
| Data pre-processing | ✗ | ✗ | ✗ | ✗ | ✓ |
| Graphical output | ✗ | ✗ | ✗ | ✗ | ✓ |
| Implementation | Matlab | Matlab script | Matlab script | Matlab toolbox | R-package Web-based app |
| Comments | The actual algorithm is not released as script or equivalent, but has to be implemented by users or used within a metabolic flux tool | Elongation calculated only for FA(18:0) | Steady state must be achieved as M+0 = import. Desaturation based on total labeling and exemplified only for FA(18:1n9). | Elongation described as de novo lipogenesis up to the total number of carbons plus multiple import-elongation terms. | |

#### Supplementary Figure legends

**Supplementary Figure 1. Chromatographic separation of the FA standards.** **a**, all the FA standards. **b**, FA(14:0). **c**, FA(14:1). **d**, FA(16:0). **e**, FA(16:1n5,n7,n9, and n10). **f**, FA(18:0). **g**, FA(18:1n7, n9, n10, and n7t). **h**, FA(18:2n6). **i**, FA(18:3n3, and n6). **j**, FA(20:0). **k**, FA(20:1n9). **l**, FA(20:2n6). **m**, FA(20:3n3, n6, and n9). **n**, FA(20:4n3, and n6). **o**, FA(20:5n3). **p**, FA(22:0). **q**, FA(22:1n9). **r**, FA(22:2n6). **s**, FA(22:3n6). **t**, FA(22:4n6). **u**, FA(22:5n3, and n6). **v**, FA(22:6n3). **w**, FA(24:0). **x**, FA(24:1n9).

**Supplementary Figure 2. Detailed workflow for data preprocessing.** Data preprocessing can be performed with a combination of our developed in-house R package LipidMS and FAMetA (Option A), or using any other suitable preprocessing tool (Option B). For data preprocessing with LipidMS, raw data files must firstly be converted into mzXML. LipidMS uses raw data files in the mzXML format and a csv metadata file as input to cover peak-peaking, alignment, grouping and peak filling. Output is a *msbatch* object that can be directly used by FAMetA to perform other preprocessing steps, including FA annotation and isotope detection. Output is a *fadata* object that can be used to conduct the final preprocessing step, which is natural abundance correction and normalisation. *Italics* depict the functions that can be executed in LipidMS or FAMetA.

**Supplementary Figure 3. Detailed FAMetA workflow and output.** Starting with the *fadata* object generated during data preprocessing (**Supplementary Figure 3**), FAMetA sequentially performs the analysis of DNS (*synthesisAnalysis* function), elongation (*elongationAnalysis* function) and desaturation (*desaturationAnalysis* function). The results for each step can be exported or a summary of all the calculated parameters and a group-based comparison can be obtained by executing the function *summarizeResults*.

**Supplementary Figure 4. *In silico* validation of the estimation of the *de novo* synthesis of FA(16:0).** To evaluate FAMetA's ability to estimate the DNS analysis parameters, realistic values for $D_1$ (5 values from 0 to 0.2), $D_2$ (15 values from 0 to 1), $\Phi$ (10 points from 0 to 0.1) and $S$ (15 values from 0 to 1) are combined to simulate 3,945 theoretical FA(16:0) distributions, to which the 0%, 2%, 5% and 10% noise levels are added to obtain 10 different noised distributions for each set of parameters. **a**, Evaluation of $S$ as a function of $S$ and $D_2$. **b**, Evaluation of $D_2$ as a function of $S$ and $D_2$. **c**, Evaluation of $S$ as a function of $S$ and $\Phi$ ($D_2$=0.5). **d**, Evaluation of $\Phi$ as a function of $S$ and $\Phi$ ($D_2$=0.5). **e**, Evaluation of $D_2$ as a function of $D_2$ and $\Phi$ ($S$ =0.5). **f**, Evaluation of $\Phi$ as a function of $D_2$ and $\Phi$ ($S$ =0.5). **g**, Evaluation of $D_1$ as a function of $D_1$ and $S$. **h**, Evaluation of $D_1$ as a function of $D_1$ and $D_2$. **i**, Evaluation of $D_1$ as a function of $D_1$ and $\Phi$.

**Supplementary Figure 5. *In silico* validation of the estimation of elongation for FA(18:0).** To evaluate FAMetA's ability to estimate parameters of elongation, the following values are set to simulate the mass-isotopologue data: $D_1$ and $\Phi$ are set at 0.05, and 0.01, respectively, $D_2$ varies from 0.1 to 0.9, and $E_1$ and $S$ from 0.05 to 1. The 0%, 25, 5% and 10% noise levels are added to obtain 10 different noised distributions for each set of parameters. **a-b**, Evaluation of $E_1$ as a function of $E_1$ and $D_2$ (**a**), and $E_1$ and $S$ (**b**). **c**, Evaluation of $S$ as a function of $E_1$ and $S$.

**Supplementary Figure 6. *In silico* validation of the estimation of elongation for FA(20:0).** To evaluate FAMetA's ability to estimate parameters of elongation, the following values are set to simulate the mass-isotopologue data: $S$, $D_1$ and $\Phi$ are set at 0.6, 0.05 and 0.01, respectively, $D_2$ varies from 0.1 to 0.9, and $E_n$ from 0.1 to 1. The 0%, 2%, 5% and 10% noise levels are added to obtain 10 different noised distributions for each set of parameters. **a-c**, Evaluation of $E_2$ as a function of $E_2$ and $D_2$ (**a**), $E_2$ and $E_1$ (**b**), and $E_2$ and $S$ (**c**). **d**, Evaluation of $E_1$ as a function of $E_2$ and $E_1$. **e**, Evaluation of $S$ as a function of $E_2$ and $S$.

**Supplementary Figure 7. *In silico* validation of the estimation of elongation for FA(22:0).** To evaluate FAMetA's ability to estimate parameters of elongation, the following values are set to simulate the mass-isotopologue data: $S$, $D_1$ and $\Phi$ are set at 0.6, 0.05 and 0.01, respectively, $D_2$ varies from 0.1 to 0.9, and $E_n$ from 0.1 to 1. The 0%, 2%, 5% and 10% noise levels are added to obtain 10 different noised distributions for each set of parameters. **a-d**, Evaluation of $E_3$ as a function of $E_3$ and $D_2$ (**a**), $E_3$ and $E_2$ (**b**), $E_3$ and $E_1$ (**c**), and $E_3$ and $S$ (**d**). **e**, Evaluation of $E_2$ as a function of $E_3$ and $E_2$. **f**, Evaluation of $E_1$ as a function of $E_3$ and $E_1$. **g**, Evaluation of $S$ as a function of $E_3$ and $S$.

**Supplementary Figure 8. *In silico* validation of the estimation of elongation for FA(24:0).** To evaluate FAMetA's ability to estimate parameters of elongation, the following values are set to simulate the mass-isotopologue data: $S$, $D_1$ and $\Phi$ are set at 0.6, 0.05 and 0.01, respectively, $D_2$ varies from 0.1 to 0.9, and $E_n$ from 0.1 to 1. The 0%, 2%, 5% and 10% noise levels are added to obtain 10 different noised distributions for each set of parameters. **a-e**, Evaluation of $E_4$ as a function of $E_4$ and $D_2$ (**a**), $E_4$ and $E_3$ (**b**), $E_4$ and $E_2$ (**c**), $E_4$ and $E_1$ (**d**), and $E_3$ and $S$ (**e**). **f**, Evaluation of $E_3$ as a function of $E_4$ and $E_3$. **g**, Evaluation of $E_2$ as a function of $E_4$ and $E_2$. **h**, Evaluation of $E_1$ as a function of $E_4$ and $E_1$. **i**, Evaluation of $S$ as a function of $E_4$ and $S$.

**Supplementary Figure 9. *In silico* validation of the estimation of desaturation.** To evaluate FAMetA's ability to estimate parameters of desaturation, the following values are set to simulate the mass-isotopologue data: $D_1$ and $\Phi$ are set at 0.05 and 0.01, respectively, $D_2$ varies from 0.1 to 0.9, $\Delta_1$ varies from 0 to 1; for FA(16:1n7), S varies from 0.1 to 1; for FA(18:1n9) S is set at 0.6 and $E_1$ varies from 0.1

to 1. The 0%, 2%, 5% and 10% noise levels are added to obtain 10 different noised distributions for each set of parameters. **a-b**, Evaluation of $\Delta_1$ for FA(16:1n7) as a function of $D_2$ and $\Delta_1$ (**a**) or $S$ and $\Delta_1$ (**b**). **c-d**, Evaluation of $\Delta_1$ for FA(18:1n9) as a function of $D_2$ and $\Delta_1$ (**c**) or $E_1$ and $\Delta_1$ (**d**).

**Supplementary Figure 10. FAMetA web application (www.fameta.es) screenshots. a**, the first tab of the FAMetA web application ("Home" tab) summarises FAMetA information. The next tabs (**b-d**) take users through the complete data processing workflow. **b**, "Data preprocessing" tab; **c**, "Manual curation" tab; **d**, "Metabolic Analysis" tab. **e**, The last tab, "Resources", contains links with access to extradocumentation, examples and the source code of the package.

#### Supplementary Figure 1

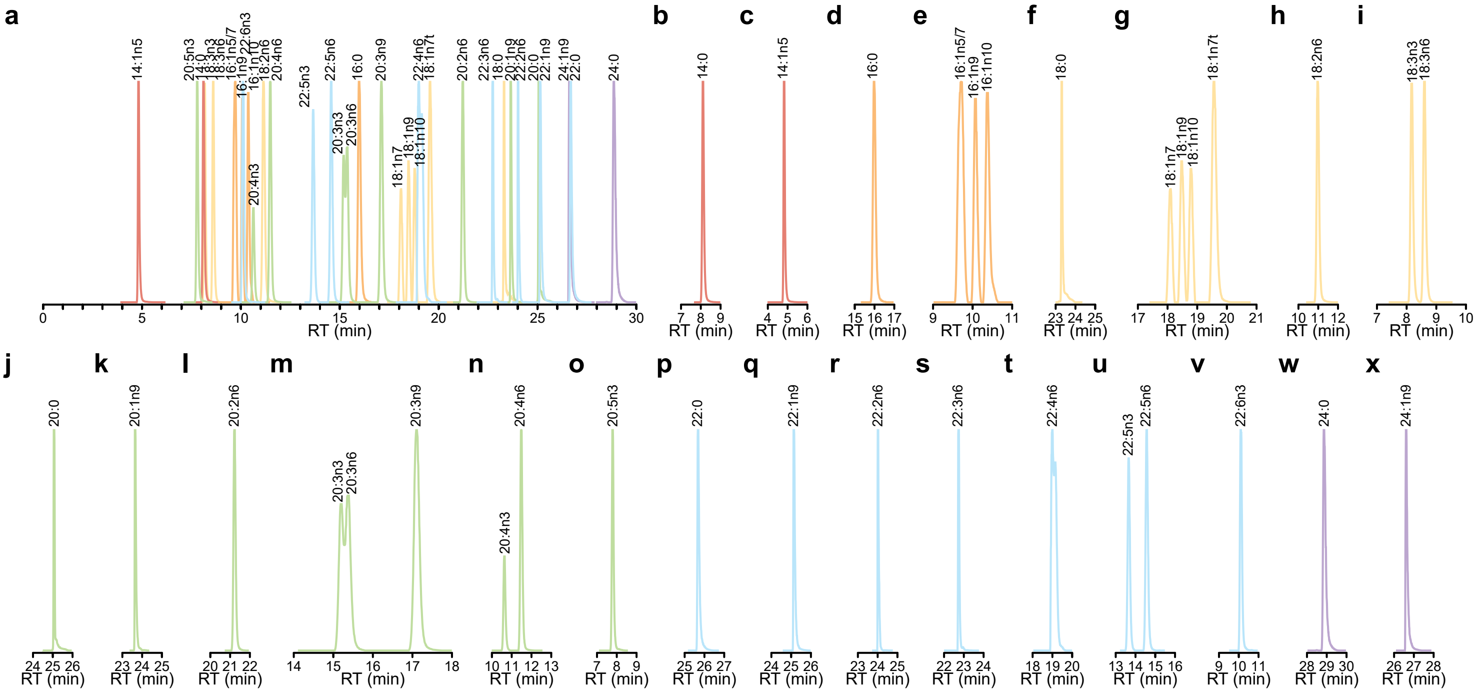

### Supplementary Figure 2

#### OPTION A

##### 1. Data conversion (MSConvert)

Convert raw data file to a format compatible with the pre-processing software. Any file converter such as MSConvert from Proteowizard may be used. In the case of LipidMS (R-package with an output directly compatible with FAMetA), mzXML format must be used.

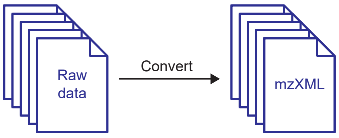


##### 2. Data pre-processing (LipidMS)

We propose LipidMS R-package to perform peak-picking, alignment and grouping. It will return an msbatch object compatible with FAMetA.

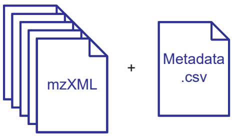


| sample | acquisitionmode | sampletype | external normalization (optional) |
|---|---|---|---|
| Sample1 | neg | class1 | 1.2 |
| Sample2 | neg | class1 | 1.1 |
| Sample3 | neg | class1 | 1.3 |
| Sample4 | neg | class2 | 1.1 |
| (…) | (…) | (…) | (…) |

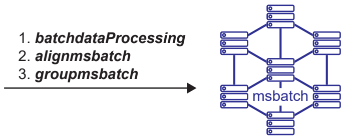


##### 3. Fatty acid annotation and manual curation

Annotate FA based on *m/z.* Annotations have to be manually curated to identify isomers.
During manual curation missing FA or internal standards can be added and parameters modified.

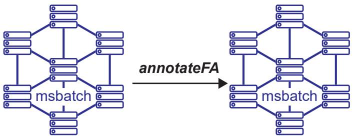


msbatch$fas

| ID | FA | *m/z* | iniRT | endRT |
|---|---|---|---|---|
| 1 | 14:0 | $m/z_1$ | $iniRT_1$ | $endRT_1$ |
| 2 | 16:0 | $m/z_2$ | $iniRT_2$ | $endRT_2$ |
| 3 | 16:1 | $m/z_3$ | $iniRT_3$ | $endRT_3$ |
| 4 | 16:1 | $m/z_4$ | $iniRT_4$ | $endRT_4$ |
| (…) | (…) | (…) | (…) | (…) |

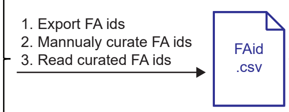


| ID | FA | *m/z* | iniRT | endRT |
|---|---|---|---|---|
| 1 | 14:0 | $m/z_1$ | $iniRT_1$ | $endRT_1$ |
| 2 | 16:0 | $m/z_2$ | $iniRT_2$ | $endRT_2$ |
| 3 | 16:1n7 | $m/z_3$ | $iniRT_3$ | $endRT_3$ |
| 4 | 16:1n10 | $m/z_4$ | $iniRT_4$ | $endRT_4$ |
| (…) | (…) | (…) | (…) | (…) |
| | IS | $m/z_{IS1}$ | $iniRT_{IS1}$ | $endRT_{IS1}$ |
| | IS | $m/z_{IS2}$ | $iniRT_{IS2}$ | $endRT_{IS2}$ |

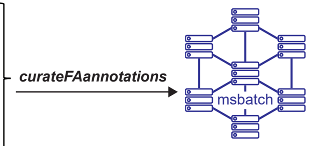


##### 4. Isotope detection

Search of $^{13}C$ isotopes based on *m/z* and peak shape correlation.

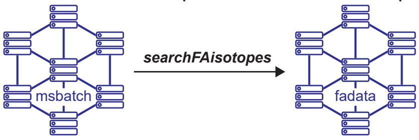


#### OPTION B

##### Import already pre-processed data (steps 1- 4)

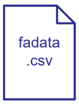


| | sampletype | class1 | class1 | class1 | class2 | (…) |
|---|---|---|---|---|---|---|
| | external normalization (opt) | 1.2 | 1.1 | 1.3 | 1.1 | (…) |
| Compound | Label | Sample1 | Sample2 | Sample3 | Sample4 | (…) |
| FA(14:0) | 0 | $I_{14:0m0}$ | $I_{14:0m0}$ | $I_{14:0m0}$ | $I_{14:0m0}$ | (…) |
| FA(14:0) | 1 | $I_{14:0m1}$ | $I_{14:0m1}$ | $I_{14:0m1}$ | $I_{14:0m1}$ | (…) |
| FA(14:0) | 2 | $I_{14:0m2}$ | $I_{14:0m2}$ | $I_{14:0m2}$ | $I_{14:0m2}$ | (…) |
| FA(14:0) | (…) | (…) | (…) | (…) | (…) | (…) |
| FA(14:0) | 14 | $I_{14:0m14}$ | $I_{14:0m14}$ | $I_{14:0m14}$ | $I_{14:0m14}$ | (…) |
| FA(16:0) | 0 | $I_{16:0m0}$ | $I_{16:0m0}$ | $I_{16:0m0}$ | $I_{16:0m0}$ | (…) |
| FA(16:0) | 1 | $I_{16:0m1}$ | $I_{16:0m1}$ | $I_{16:0m1}$ | $I_{16:0m1}$ | (…) |
| FA(16:0) | 2 | $I_{16:0m2}$ | $I_{16:0m2}$ | $I_{16:0m2}$ | $I_{16:0m2}$ | (…) |
| FA(16:0) | (…) | (…) | (…) | (…) | (…) | (…) |
| FA(16:0) | 16 | $I_{16:0m16}$ | $I_{16:0m16}$ | $I_{16:0m16}$ | $I_{16:0m16}$ | (…) |
| (…) | (…) | (…) | (…) | (…) | (…) | (…) |

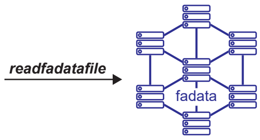


##### 5. Natural abundance correction and normalization

Correct for natural abundance of $^{13}C$ isotopes using the Accucor algorithm and, if available, IS-based normalization, blank substraction and  external factor normalization (e.g. protein, cell number...).

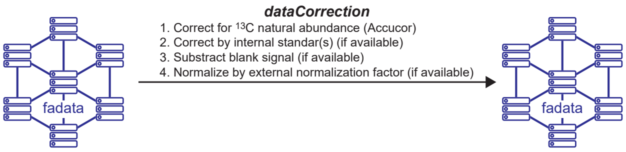


fadata$intensities

| Compound | Label | Sample1 | Sample2 | Sample3 | Sample4 | (…) |
|---|---|---|---|---|---|---|
| FA(14:0) | 0 | $I_{14:0m0}$ | $I_{14:0m0}$ | $I_{14:0m0}$ | $I_{14:0m0}$ | (…) |
| | 1 | $I_{14:0m1}$ | $I_{14:0m1}$ | $I_{14:0m1}$ | $I_{14:0m1}$ | (…) |
| | 2 | $I_{14:0m2}$ | $I_{14:0m2}$ | $I_{14:0m2}$ | $I_{14:0m2}$ | (…) |
| | (…) | (…) | (…) | (…) | (…) | (…) |
| | 14 | $I_{14:0m14}$ | $I_{14:0m14}$ | $I_{14:0m14}$ | $I_{14:0m14}$ | (…) |
| FA(16:0) | 0 | $I_{16:0m0}$ | $I_{16:0m0}$ | $I_{16:0m0}$ | $I_{16:0m0}$ | (…) |
| | 1 | $I_{16:0m1}$ | $I_{16:0m1}$ | $I_{16:0m1}$ | $I_{16:0m1}$ | (…) |
| | 2 | $I_{16:0m2}$ | $I_{16:0m2}$ | $I_{16:0m2}$ | $I_{16:0m2}$ | (…) |
| | (…) | (…) | (…) | (…) | (…) | (…) |
| | 16 | $I_{16:0m16}$ | $I_{16:0m16}$ | $I_{16:0m16}$ | $I_{16:0m16}$ | (…) |
| (…) | (…) | (…) | (…) | (…) | (…) | (…) |

fadata$poolsize
fadata$relativepoolsize

| Sample | Sample1 | Sample2 | Sample3 | Sample4 | (…) |
|---|---|---|---|---|---|
| Class | Class1 | Class1 | Class1 | Class2 | (…) |
| FA(14:0) | $I_{14:0}$ | $I_{14:0}$ | $I_{14:0}$ | $I_{14:0}$ | (…) |
| FA(14:1n5) | $I_{14:1n5}$ | $I_{14:1n5}$ | $I_{14:1n5}$ | $I_{14:1n5}$ | (…) |
| FA(16:0) | $I_{16:0}$ | $I_{16:0}$ | $I_{16:0}$ | $I_{16:0}$ | (…) |
| FA(16:1n7) | $I_{16:1n7}$ | $I_{16:1n7}$ | $I_{16:1n7}$ | $I_{16:1n7}$ | (…) |
| FA(16:1n9) | $I_{16:1n9}$ | $I_{16:1n9}$ | $I_{16:1n9}$ | $I_{16:1n9}$ | (…) |
| FA(16:1n10) | $I_{16:1n10}$ | $I_{16:1n10}$ | $I_{16:1n10}$ | $I_{16:1n10}$ | (…) |
| FA(18:0) | $I_{18:0}$ | $I_{18:0}$ | $I_{18:0}$ | $I_{18:0}$ | (…) |
| (…) | (…) | (…) | (…) | (…) | (…) |

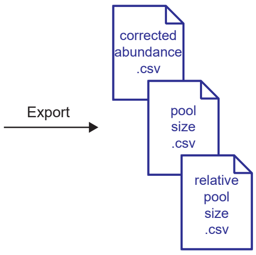

### Supplementary Figure 3

#### 1. Analysis of de novo syntesis (carbon number ≤16)

Estimate fraction of newly synthesized FA and contribution of the $^{13}C$ tracer to acetate pool using a quasi-multinomial distribution.

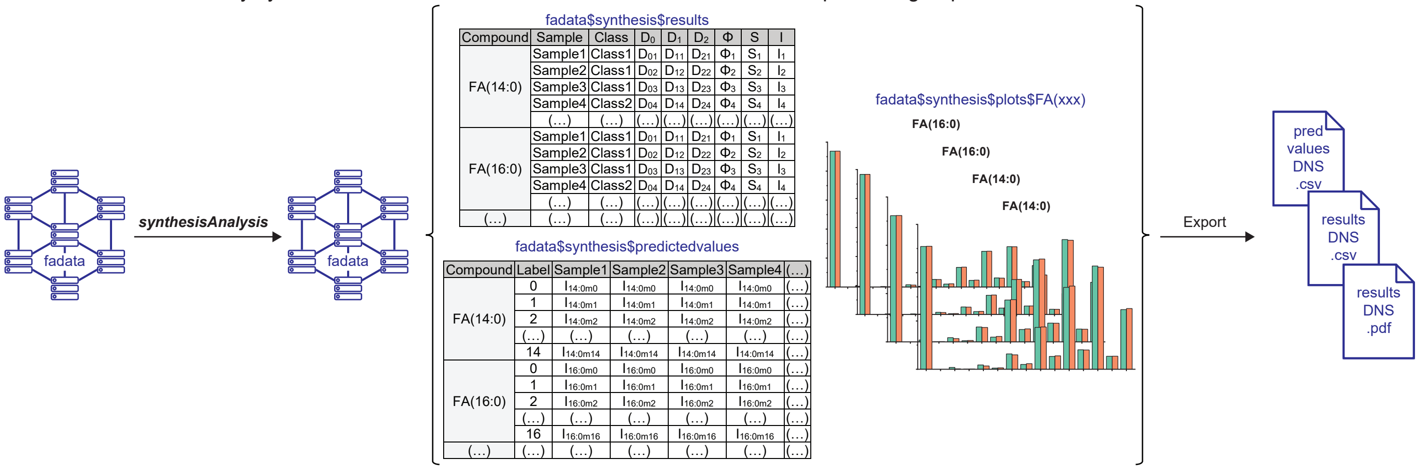


#### 2. Analysis of elongation

Estimate fraction of elongated FA.

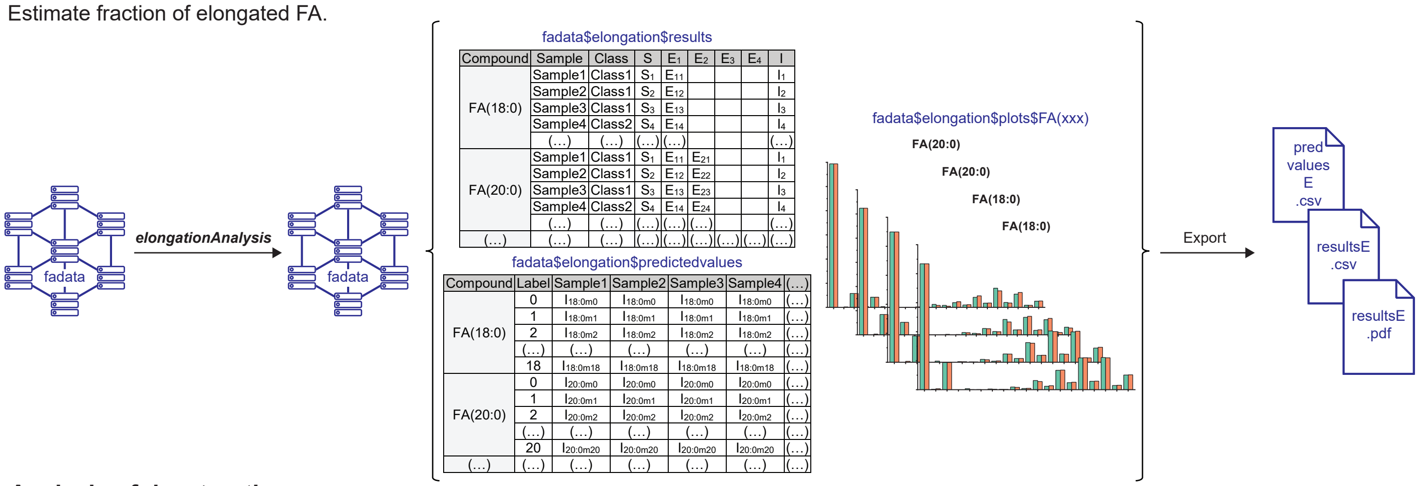


#### 3. Analysis of desaturation

Estimate the fraction of desaturated FA based on previous results for synthesis and elongation analysis.

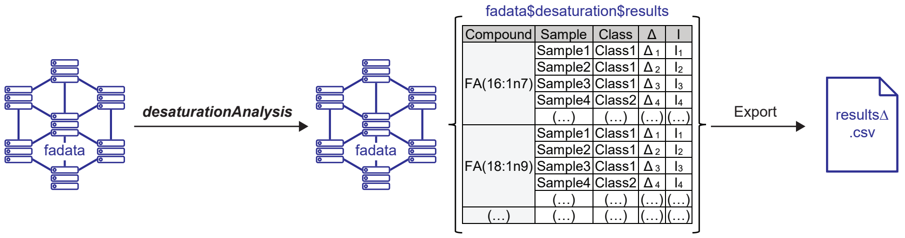


#### 4. Summary and graphical output

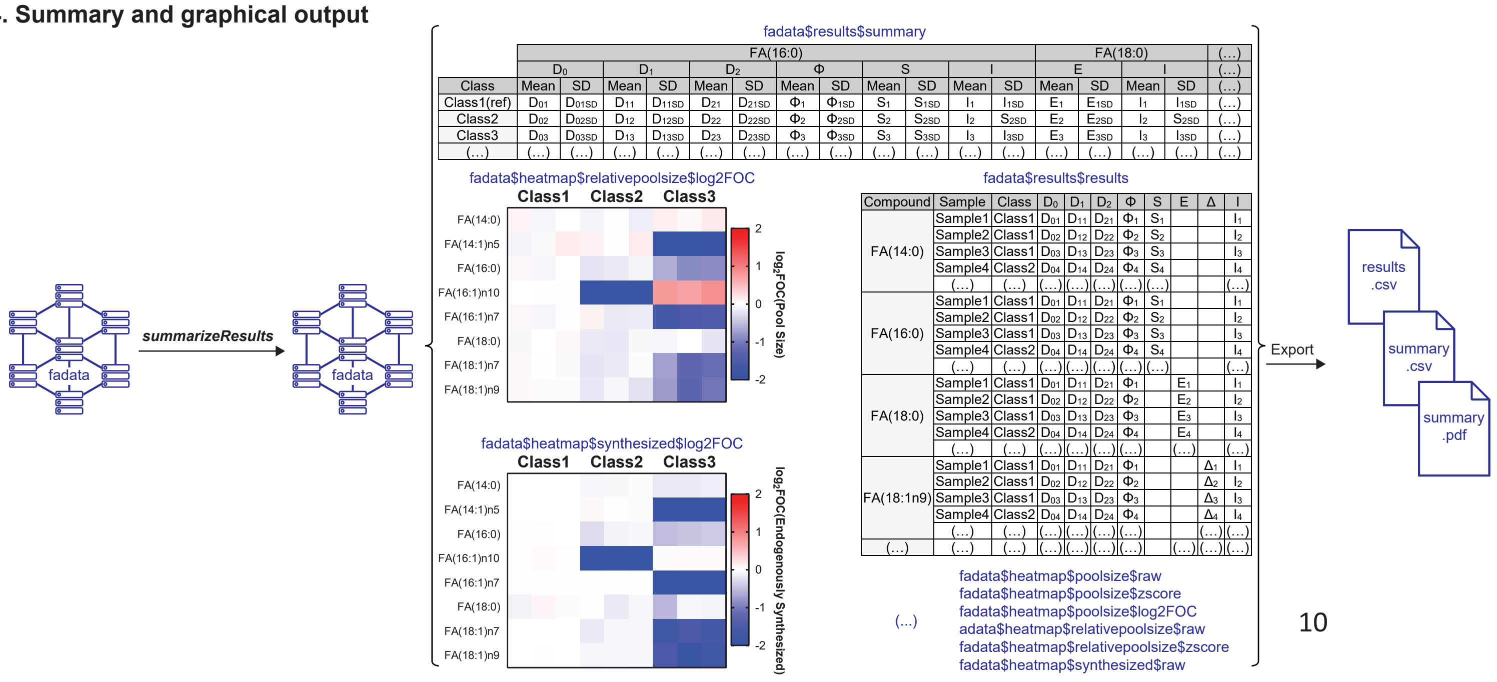

### Supplementary Figure 4

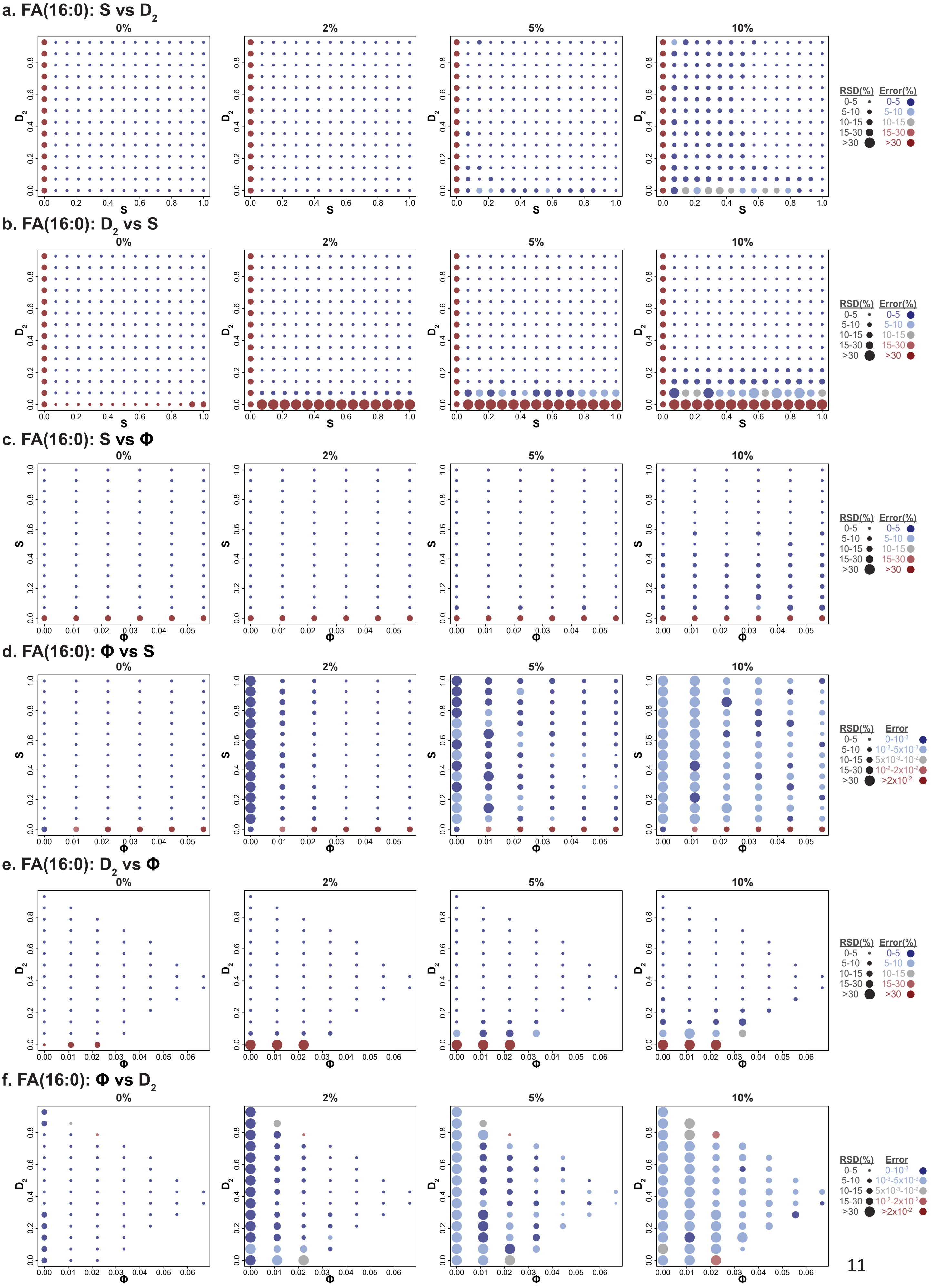

### Supplementary Figure 4 (continued)

## g. FA(16:0): $D_1$ vs S

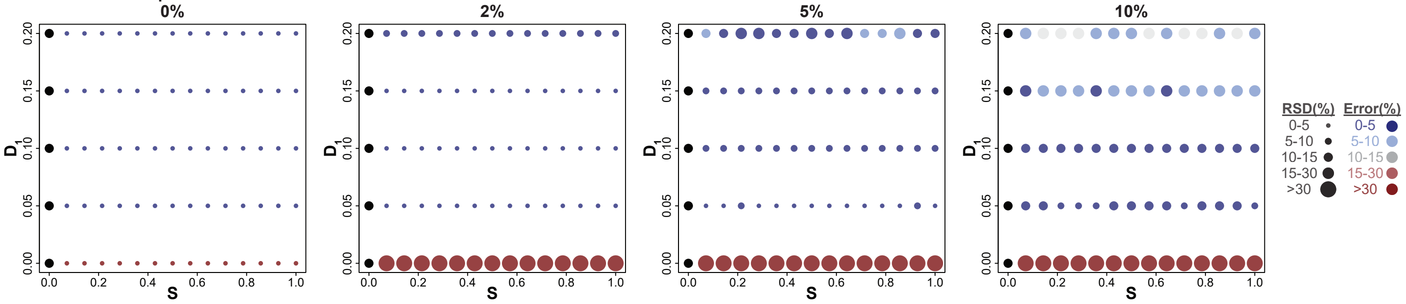


## h. FA(16:0): $D_1$ vs $D_2$

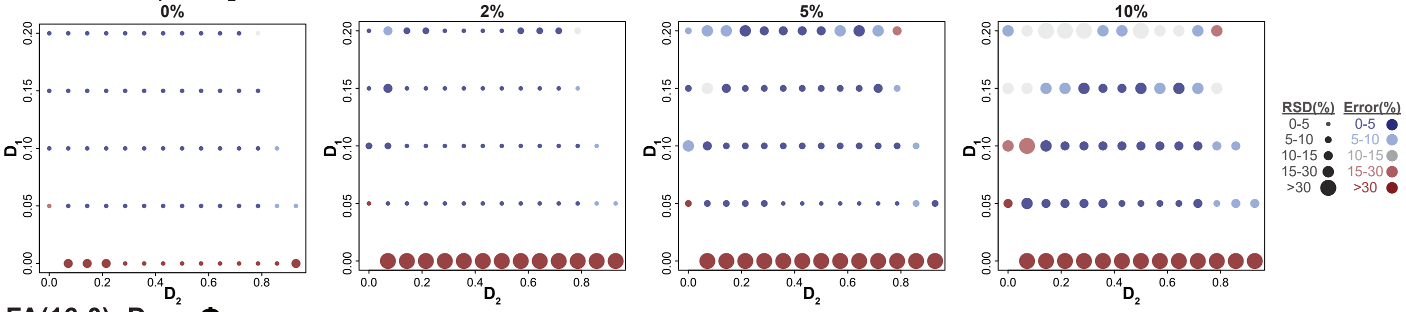


## i. FA(16:0): $D_1$ vs Φ

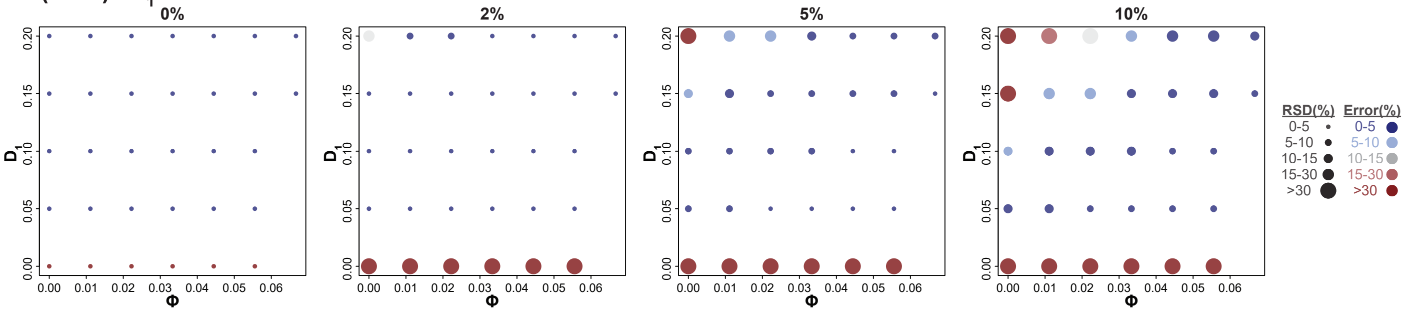

### Supplementary Figure 5

## a. FA(18:0): $E_1$ vs $D_2$

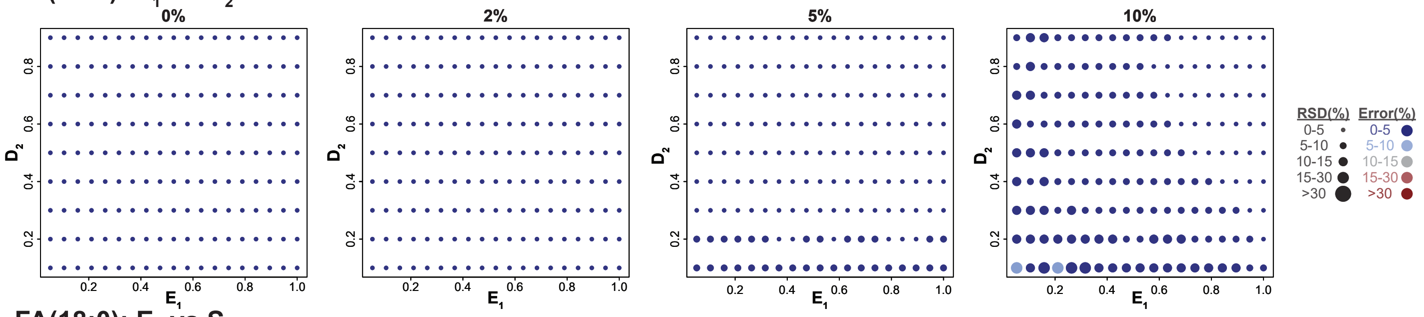


## b. FA(18:0): $E_1$ vs S

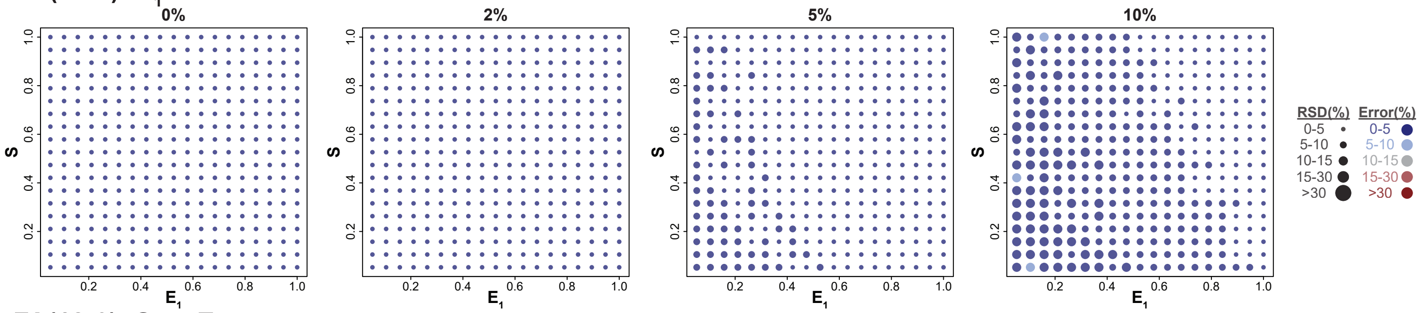


## c. FA(18:0): S vs $E_1$

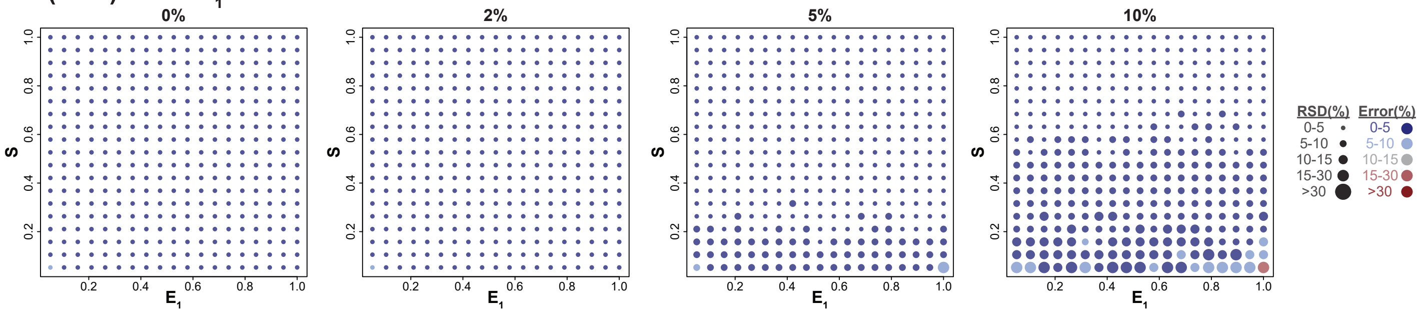

### Supplementary Figure 6

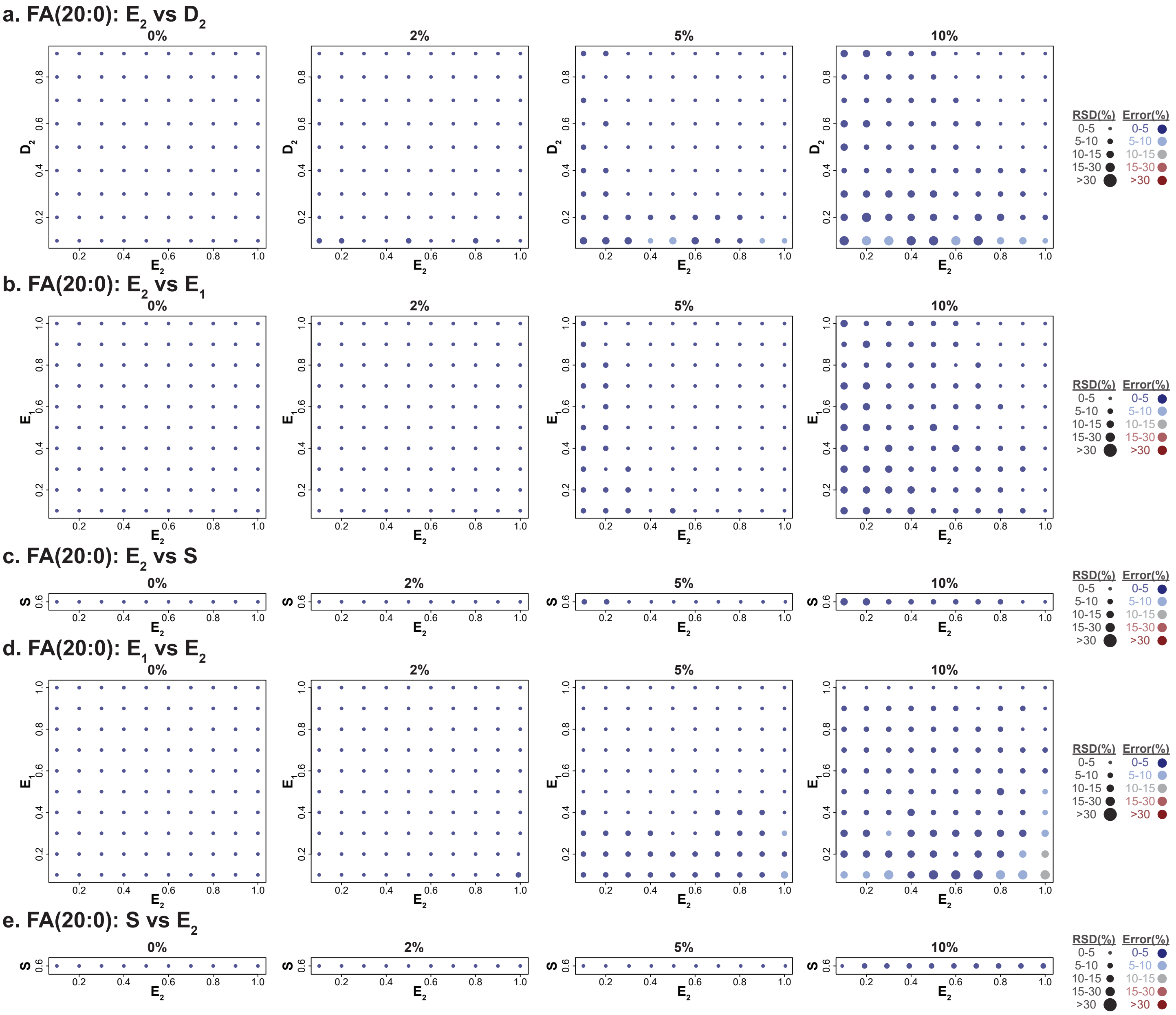

#### Supplementary Figure 7

### a. FA(22:0): $E_3$ vs $D_2$

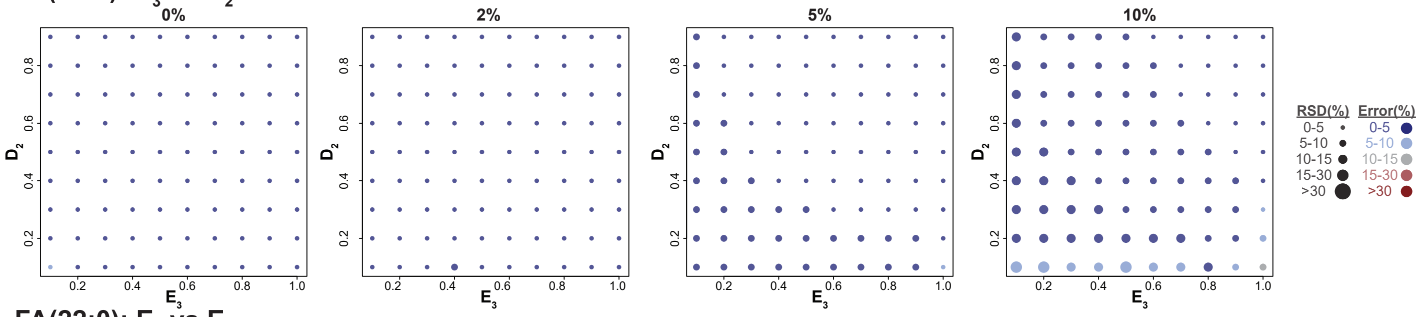


### b. FA(22:0): $E_3$ vs $E_2$

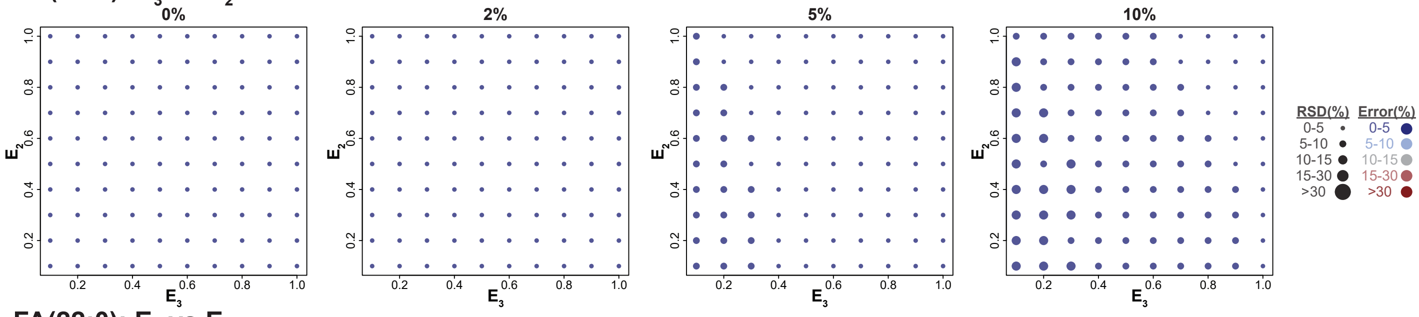


### c. FA(22:0): $E_3$ vs $E_1$

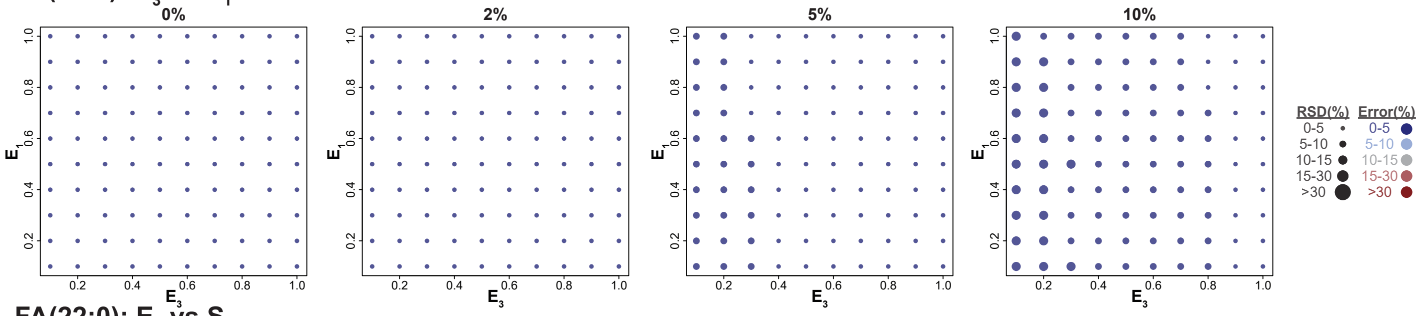


### d. FA(22:0): $E_3$ vs S

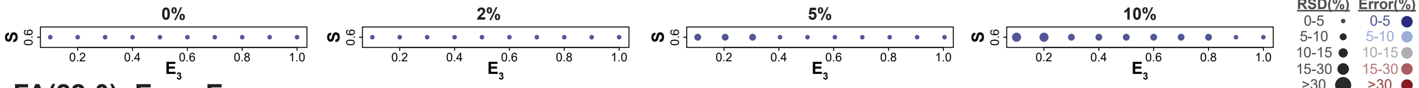


### e. FA(22:0): $E_2$ vs $E_3$

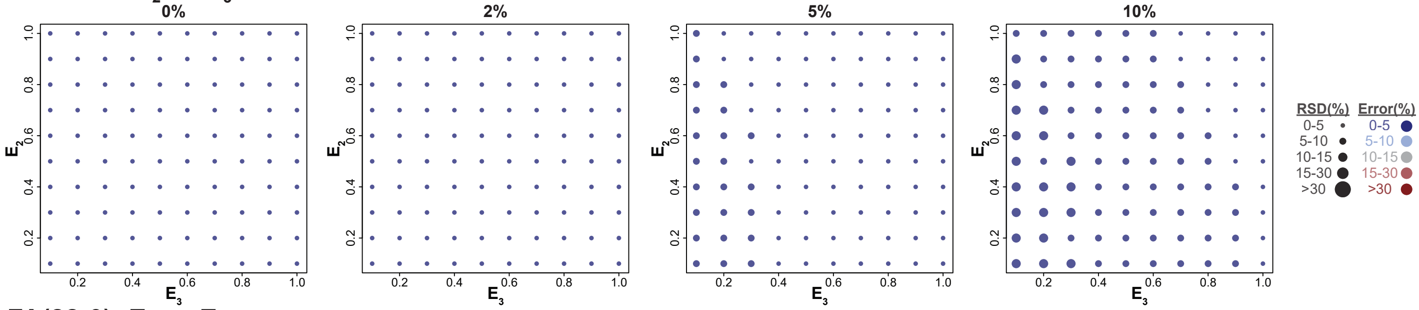


### f. FA(22:0): $E_1$ vs $E_3$

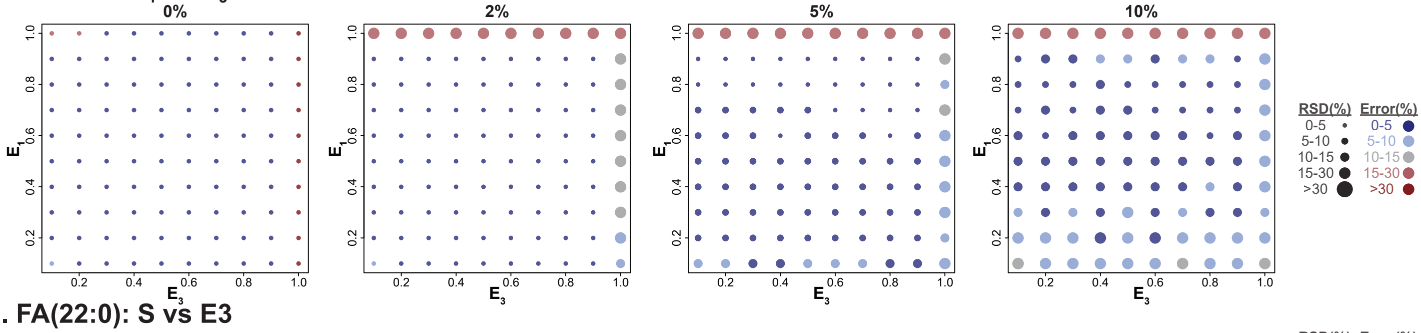


### g. FA(22:0): S vs E3

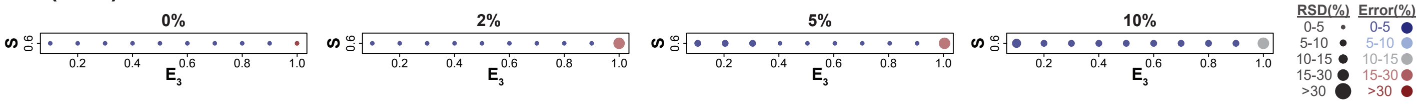

#### Supplementary Figure 8

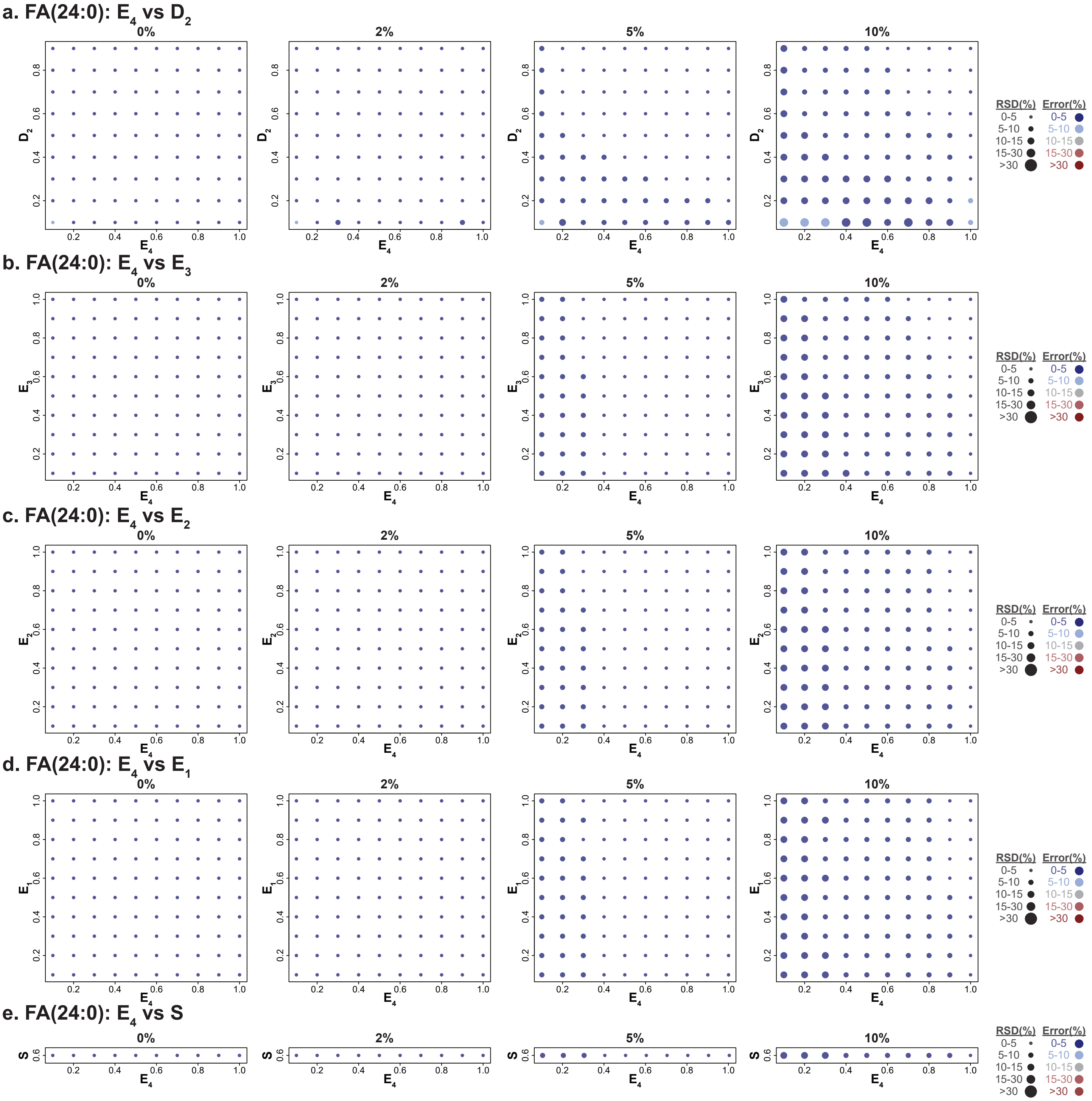

### Supplementary Figure 8 (continued)

f. FA(24:0): $E_3$ vs $E_4$

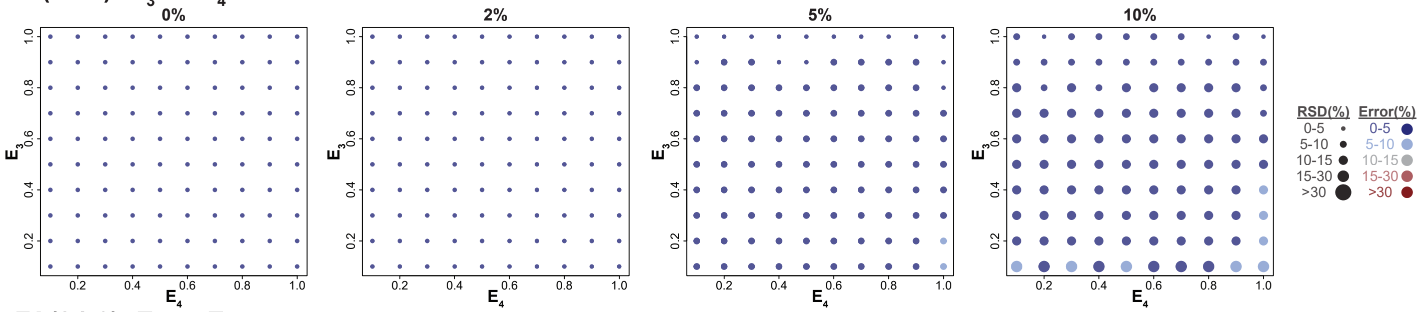


g. FA(24:0): $E_2$ vs $E_4$

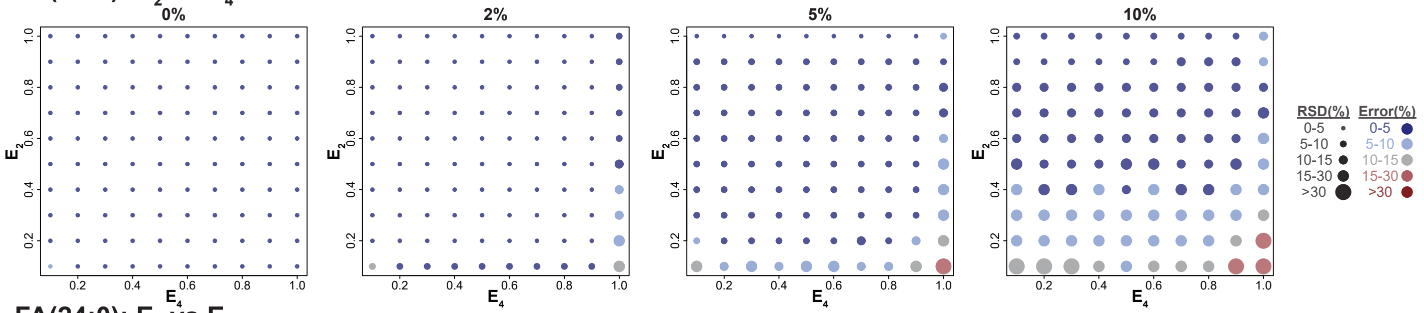


h. FA(24:0): $E_1$ vs $E_4$

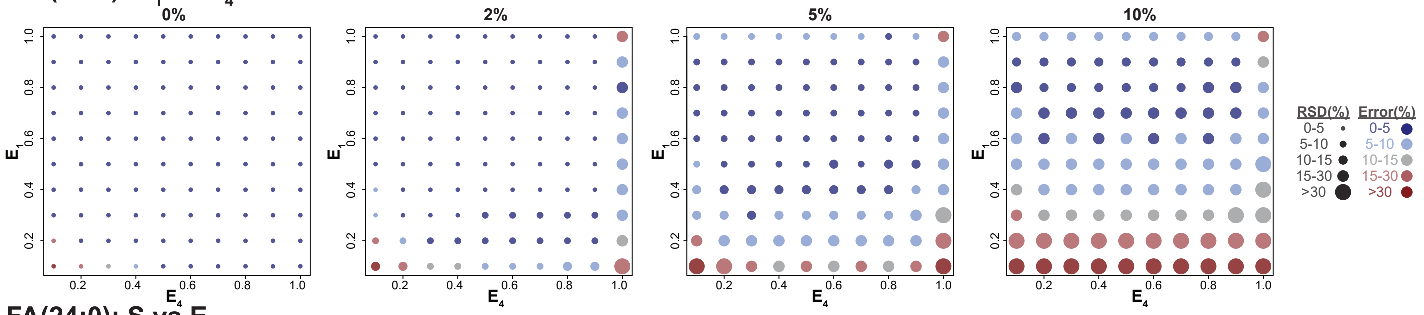


i. FA(24:0): S vs $E_4$

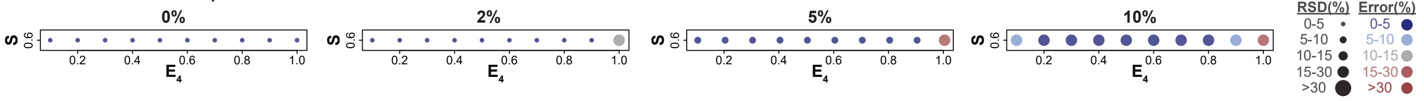

### Supplementary Figure 9

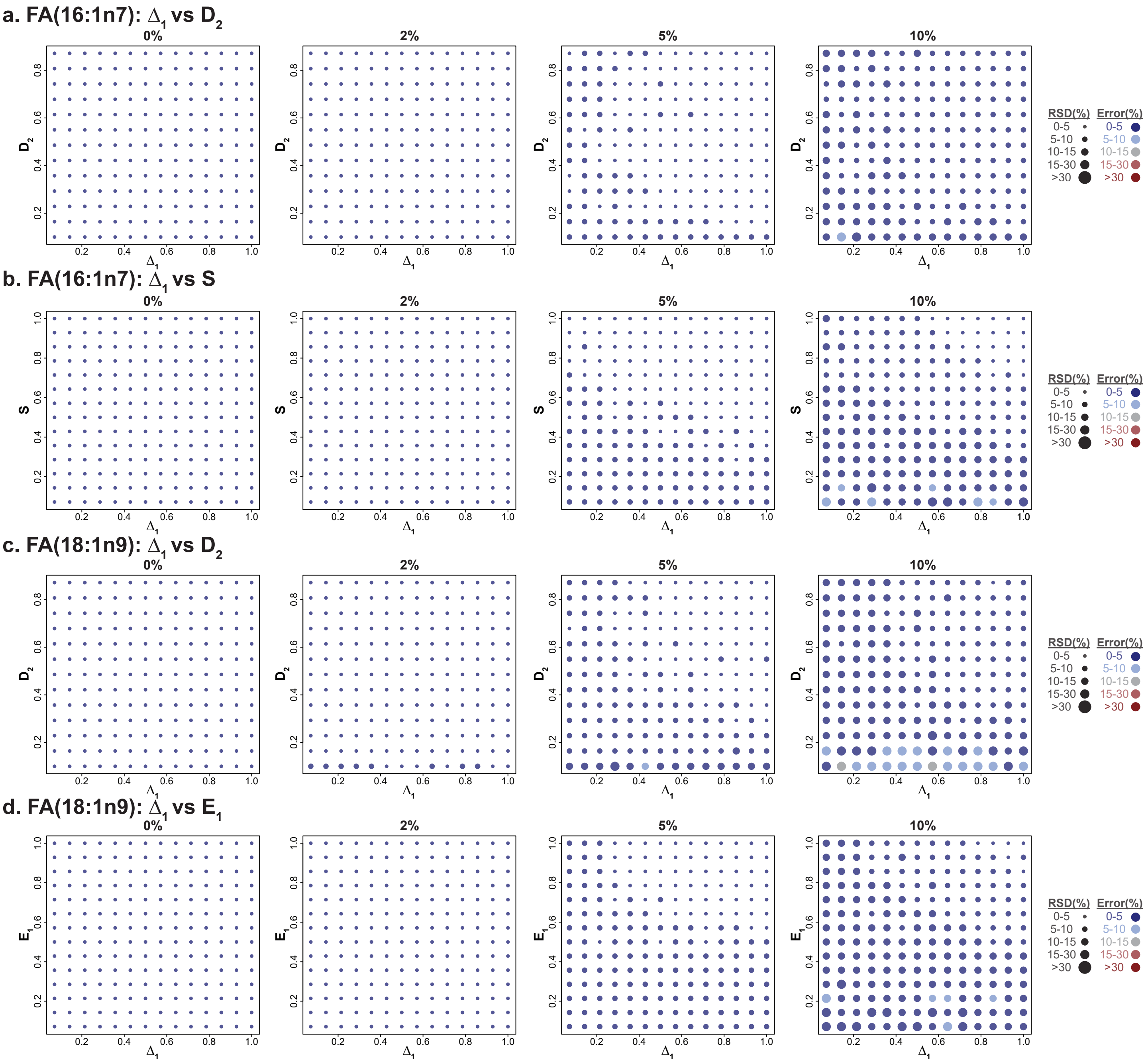

#### Supplementary Figure 10

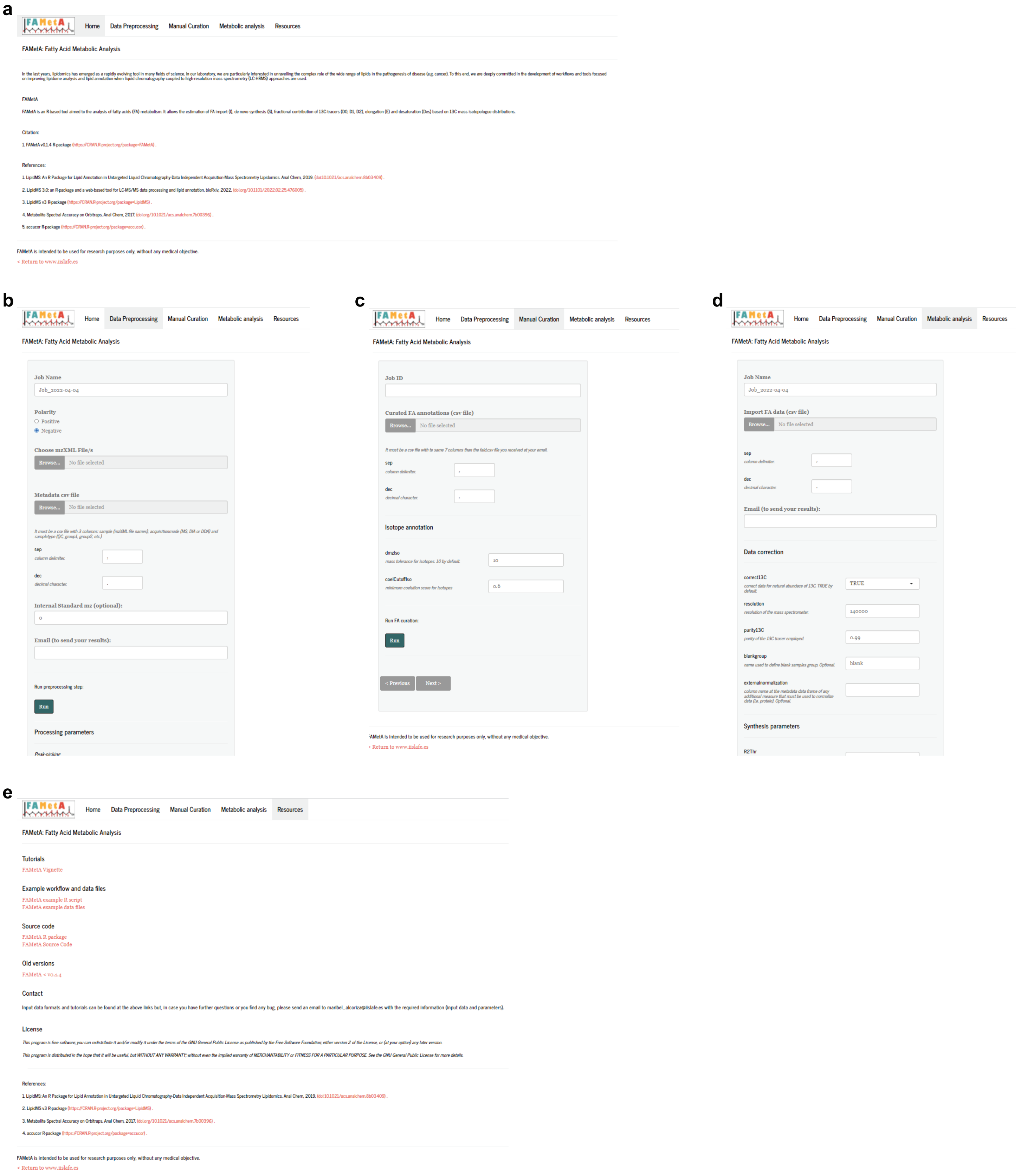
